# Infectious disease phylodynamics with occurrence data

**DOI:** 10.1101/596700

**Authors:** Leo A. Featherstone, Francesca Di Giallonardo, Edward C. Holmes, Timothy G. Vaughan, Sebastián Duchêne

## Abstract

**Point 1:** Phylodynamic models use pathogen genome sequence data to infer epidemiological dynamics. With the increasing genomic surveillance of pathogens, especially amid the SARS-CoV-2 outbreak, new practical questions about their use are emerging.

**Point 2:** One such question focuses on the inclusion of un-sequenced case occurrence data alongside sequenced data to improve phylodynamic analyses. This approach can be particularly valuable if sequencing efforts vary over time.

**Point 3:** Using simulations, we demonstrate that birth-death phylodynamic models can employ occurrence data to eliminate bias in estimates of the basic reproductive number due to misspecification of the sampling process. In contrast, the coalescent exponential model is robust to such sampling biases, but in the absence of a sampling model it cannot exploit occurrence data. Subsequent analysis of the SARS-CoV-2 epidemic in the northwest USA supports these results.

**Point 4:** We conclude that occurrence data are a valuable source of information in combination with birth-death models. These data should be used to bolster phylodynamic analyses of infectious diseases and other rapidly spreading species in the future.

## Introduction

Outbreak investigations increasingly rely on genome sequencing of causative pathogens. Phylodynamic methods take advantage of these data to infer epidemiological dynamics (Rife et al., 2017). New sequencing technologies generate these data rapidly, such that phylodynamic inferences can be conducted in actionable time frames (Gardy & Loman, 2018; Grubaugh et al., 2019; Hadfield et al., 2018). In this context, the main appeal of phylodynamics is that it uses sequence data to infer epidemiological dynamics preceding the earliest collected sample, or during periods without collected sequences, and offers insight into transmission chains.

Phylodynamic models describe a branching process, modelling both how a branching transmission chain and phylogenetic tree of the underlying pathogen evolve. These are central to linking epidemiological dynamics to the evolution of a pathogen. In Bayesian phylogenetic implementations the particular model of a branching process is part of the prior and is sometimes referred to as the ‘tree prior’, such as the birth-death or coalescent exponential. Internal nodes in the tree are associated with transmission events while the tips of the tree represent sampling events (du Plessis & Stadler, 2015). The basic reproductive number, *R_0_*, is a key parameter that reflects the average number of secondary infections in a fully susceptible population. The simplest tree priors that can infer *R_0_* posit that the number of infected individuals increases exponentially over time. Although more sophisticated methods now exist (Kühnert et al., 2014; Popinga et al., 2015; Rasmussen et al., 2017; Vaughan et al., 2019; Volz & Siveroni, 2018), we focus here on tree priors assuming simple exponential growth since they are appropriate for the early stages of an outbreak and are increasingly used to assess the efficacy of public health interventions (Geoghegan et al., 2020; Vasylyeva et al., 2019).

Two commonly used phylodynamic tree priors are the coalescent exponential and the birth-death, both of which assume that the infected population size, *N*, grows at a rate *r*; *N(t)=e^rt^*, where *t* is time after the origin. From an epidemiological perspective, *r* is the difference between the transmission rate, *λ*, and the become uninfectious rate, *δ*, (*r*= *λ* - *δ*). 1/*δ* is the duration of infection. *R_0_* is estimated as *R_0_= λ/δ*. The exponential coalescent is a generalisation of the Kingman-n coalescent where population size is a deterministic function of time (Griffiths & Tavare, 1994; Volz et al., 2009, 2013). In contrast, the birth-death tree prior assumes stochastic population growth with sampling through time (Stadler, 2010; Stadler et al., 2012; Stadler & Yang, 2013). This is captured in the death rate*δ=ψ+μ*, where *μ* is the recovery rate and *ψ* is the sampling rate such that the sampling proportion, *p*, can be calculated as 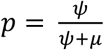.

Phylodynamic analyses draw from sequence data and sampling times (Biek et al., 2015; Drummond et al., 2002, 2003; Rambaut, 2000; Rieux & Balloux, 2016). In the coalescent exponential, sampling times are useful insofar as they influence the distribution of coalescent events through time, influencing *R_0_* in turn. Coalescent models typically condition upon sampling times instead of using them to infer sampling rates. Some ‘augmented likelihood’ approaches can combine the coalescent with a sampling process (Volz & Frost, 2014), but they are not standard practice. For the birth-death tree prior, the number of samples and their times are naturally informative because they are explicitly modelled through the sampling rate (i.e. they inform *ψ*) (Boskova et al., 2018). This is a well understood difference between the two tree priors, but its consequences remain to be explored in the context of occurrence data. Although the amount of sequence data in outbreak investigations has increased, a key consideration is that sequencing efforts are often conducted only after relatively a large number of cases are reported. This latency in sampling can bias estimates of epidemiological parameters. To visualise this, the trees in Fig 1 were simulated under an *R_0_* of 2, a constant sampling effort, and over the course of 1 year. If sequencing were only conducted for samples collected after 0.75 years, samples from the deep sections of the tree would be missed (*late sampling* in Fig 1). Such sampling bias can mislead inferences of epidemiological dynamics because there is no sampling data and very few branching events to inform inferences of the early stages of the outbreak.

**Fig 1.**
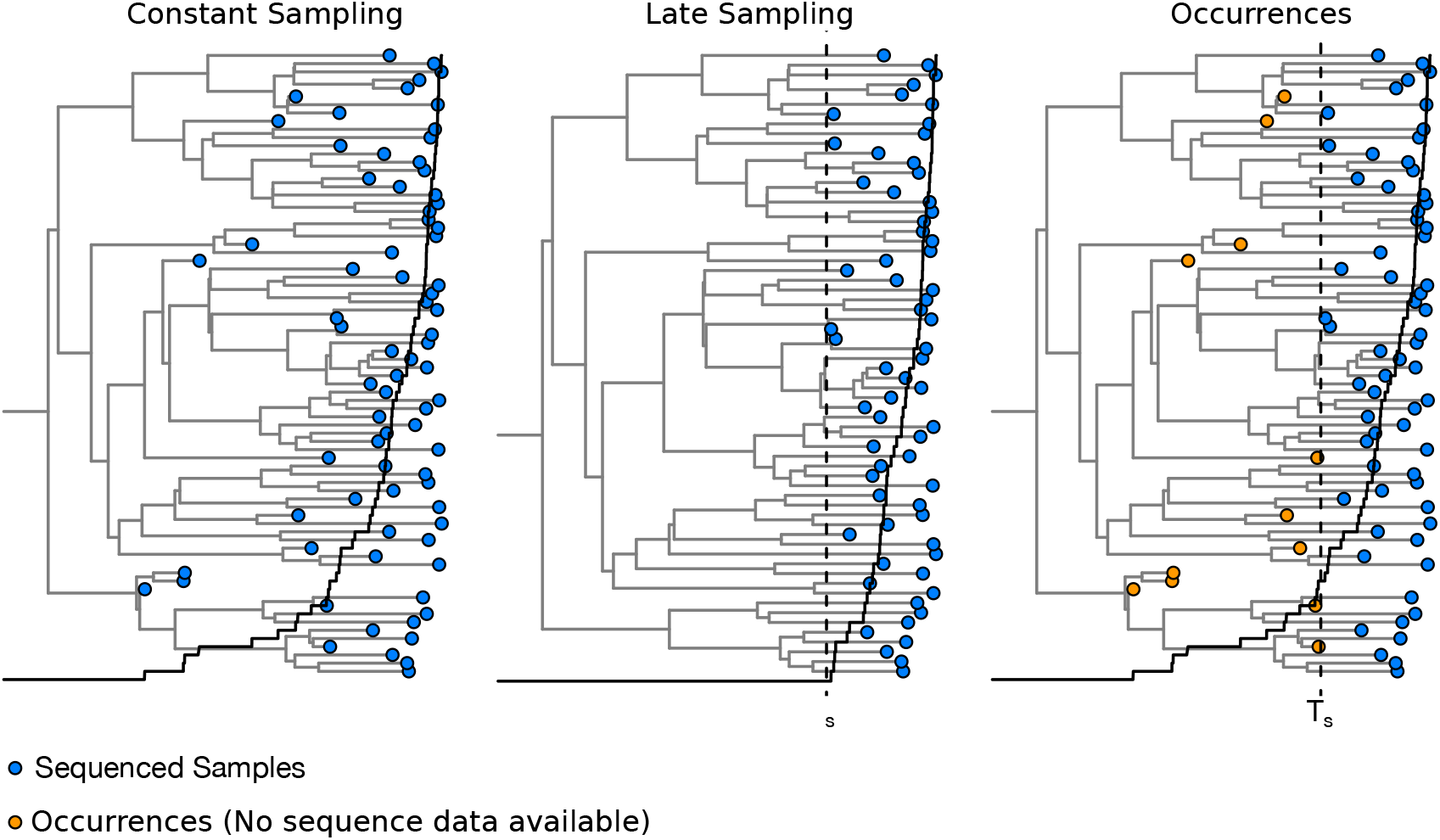
Example of a phylogenetic trees generated under a birth-death process with a basic reproductive number (*R_0_*) of 2, and a becoming uninfectious rate(*δ*) of 100 for three analysis scenarios. The solid line denotes the number of samples collected over time. In *constant* sampling samples are collected and sequenced at a rate *ψ*=5 (i.e. sampling probability, *p*, of 0.05). In *late sampling* samples are collected and sequenced after time T_s_ shown with the dashed line. In *occurrence data* samples are collected constantly over time, but only sequenced after time T_s_, such that before T_s_ only occurrences (sampling times with no sequence data) are included. Blue circles represent samples with sequence data, whereas those in orange correspond to occurrences. In the *occurrence data* scenario, a Bayesian analysis would integrate over their phylogenetic uncertainty. The solid line represents the number of samples collected over time. In *late sampling* there are no samples collected before T_s_, such that assuming constant sampling can produce a bias in estimates of epidemiological dynamics.

Here we investigate bias in epidemiological parameters due to sampling heterogeneity and present two approaches to reduce such bias using occurrence data. The first approach involves using a birth-death skyline tree prior that requires an understanding of the sampling effort (Stadler et al., 2013). If it is known that there was no attempt to collect samples early in the outbreak, one can set two intervals for the *ψ* parameter where one is zero. However, without knowledge of sampling effort this scenario is indistinguishable from a constant sampling effort where initial prevalence was so low as to preclude obtaining any sequence data early in an outbreak. The second approach consists of including early case occurrences in analyses, where an occurrence is a laboratory confirmed case that was not sequenced (*occurrences* scenario in Fig 1). Occurrence data are a relatively inexpensive and often readily available source of information because they are traditionally used in epidemiology and accurately identified via contact tracing and testing efforts. In a Bayesian phylogenetic framework, topological uncertainty due to occurrence data is naturally incorporated into the analysis through the posterior. An analogous approach can be used to coherently specify fossil data for molecular clock calibration (Heath et al., 2014; Heath & Moore, 2014). This approach and others have been modelled, but not applied in phylodynamics hitherto (Gupta et al., 2020; Manceau et al., 2019).

## Materials and Methods

### Simulation study

We simulated phylogenetic trees under a birth-death process in MASTER v6.1 (Vaughan & Drummond, 2013), with the following parameterisation; *R_0_*=2 or 1.5, *δ*=91, *p*=0.05, and an outbreak duration of one year (1/*δ* = 0.011 years, corresponding to an expected infectious period of about 4 days). The number of tips and their ages are naturally variable (from 100 to 150 tips). We assumed a strict molecular clock with an evolutionary rate of 0.01 substitutions per site per year (subs/site/year) and the HKY+Γ substitution model to produce alignments of 13,000 nucleotides using NELSI (Ho et al., 2015) and Phangorn v2.4 (Schliep, 2011). These settings are broadly similar to an influenza virus outbreak (Hedge et al., 2013), but a rescaling of the epidemiological parameters could apply to many other pathogens. We then assumed three sampling scenarios: (i) *constant* sampling with all sequences from the simulation included (e.g. the sequence for every sample in the tree in Fig 1 is included), (ii) *late sampling* only with samples after time T_s_ (e.g. only sequences for samples after the dashed line in the tree in Fig 1), and (iii) *occurrences* in which sequence data are available only after time T_s_ with those preceding recorded as occurrences. We set T_s_ to 0.75 or 0.9 years. For each parameter configuration we simulated 100 sequence data sets which were subsampled according to the three scenarios above. Occurrences were emulated by replacing simulated DNA sequences with ‘n’ (i.e. missing data) in the alignment. We analysed the data in BEAST v2.5 (Bouckaert et al., 2019) with coalescent exponential and the birth-death tree priors. Our results focus on the birth-death, but the coalescent exponential forms a valuable point of comparison through its robustness to variation in sampling. For the *late sampling* scenario, we also considered the birth-death skyline (BDSky in figures) with two intervals for the *ψ* parameter, with the interval time fixed at T_s_. We matched the substitution and clock model to those used to generate the data and we used an informative prior on *δ* using a Γ distribution with mean fixed to the true value of 91 and standard deviation of 1.

We assessed the effectiveness of each analysis treatment using three statistics. First, we considered the coverage as a measure of accuracy, or the number of times the 95% highest posterior density (HPD) intervals covered the true value of a given parameter. Second, we consider ‘average bias’, which is the difference between the posterior mean and true mean for a given parameter averaged across the 100 simulations for each sampling treatment. Third, we consider average 95% HPD width for each treatment, as a measure of precision.

### Empirical case study

To illustrate the accuracy of occurrence data relative to completely sequenced data sets we analysed 821 whole genome sequences sampled from the SARS-CoV-2 pandemic from Washington State, USA, and the adjacent Washington County, Oregon, downloaded from GISAID (Supplementary material) and partially documented by (Bedford et al., 2020). Accordingly, we downloaded 2,164 high-coverage genome sequences collected between January 18 and June 30 2020, but selected the 821 sequences taken up to March 21 2020 to capture an exponential phase in the epidemic and sampling (Fig S1). We corroborated exponential growth in the underlying population using an Epoch Sampling Proportion Skyline Plot (Parag et al., 2020). We further divided this data set into five subsets as per our simulation study: (i) ‘complete sampling’ including all 821 sequences; (ii) late sampling post March 6 2020 (decimal date 2020.18) including 637 sequences; (iii) late sampling post March 14 2020 (2020.20) including 340 sequences; (iv) late sampling post March 6 2020 (2020.18) including 637 sequences and 184 occurrences; and (v) late sampling post 2020.2 including 340 sequences and 481 occurrences. Including two late sampling data subsets offers information about how inflation in *R_0_* varies with latency in sequences.

We then analysed each data set with each tree prior used in the simulation study with BEASTv2.5. We first employed a birth-death model with serial sampling. We placed a lognormal prior on *R_0_* with mean 0 and standard deviation of 1; fixed *δ* at 36.5 (i.e. 10-day duration of infection as estimated recently (Price et al., 2020)); a β prior on sampling proportion with shape and scale equal to 2 to penalise extreme values. Second, we used a birth-death skyline with the same priors as the birth-death, but with two sampling rate parameters. The first pertained to after the 2020.18 or 2020.2 cut-off, and the second to before the cut-off. Both used the same beta prior for sampling proportion as for the birth-death. Third, a coalescent exponential tree prior was used with a Laplace prior on growth rate with mean 0 and scale 100 and an exponential prior with mean 100 on the coalescent exponential effective population size (*ϕ*). For both tree priors, we assumed HKY+Γ substitution model with a strict molecular clock rate fixed to 10^-3^ subs/site/year, following recent estimates (Duchene et al., 2020). We ran a Markov chain Monte Carlo of 5×10^8^ steps, sampling at every 1000^th^ step. We determined sufficient sampling from the posterior by verifying that the effective sample size all parameters of interest was above 200.

## Results

### Simulation study

Analyses of data sets with late sampling using the birth-death model were least accurate in estimating *R_0_*. In only 12 of 100 simulations with *R_0_*=2 did the 95% HPD include 2 (Table 1 and Fig 2a). The true value was never recovered for simulations with *R_0_*=1.5 (Table S1 and Fig S2). The birth-death skyline was more accurate with 95 and 92 of 100 simulations covering *R_0_*=2 and *R_0_*=1.5 respectively. The coalescent exponential was also more accurate with 100 and 80 simulations having HPD intervals that covered *R_0_*=2 and *R_0_*=1.5 respectively. However, this came at the cost of low precision as HPD width was the largest for the coalescent out of all treatments.

**Table 1.**
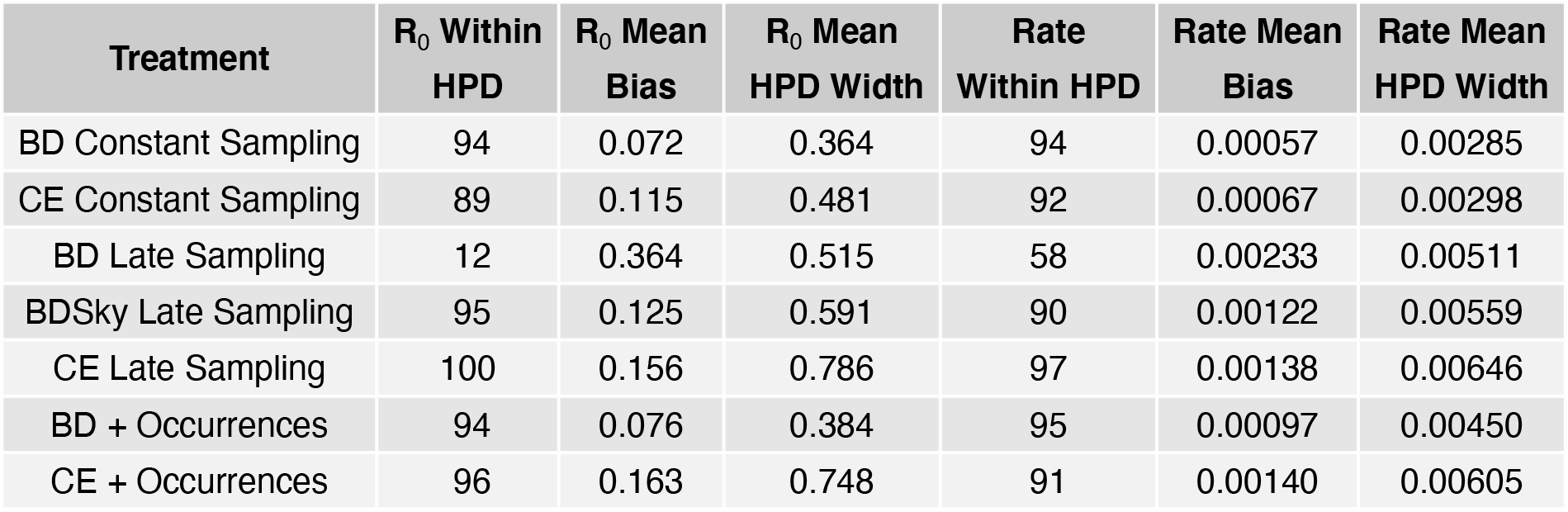
Results of the simulation study with *R_0_* of 2 and evolutionary rate of 0.01 subs/site/year. The rows correspond to the seven treatments. For *R_0_* and evolutionary rate (subs/site/year), columns denote the number of simulations (out of 100) where the value used to generate the data was contained within the 95% highest posterior density (HPD), also referred to as coverage and reflecting accuracy; average bias measured the average difference between posterior mean *R_0_* and 2; and the average HPD width. BD stands for birth-death, CE for coalescent exponential, and BDSky to the birth-death skyline model with two sampling intervals.

**Fig 2.**
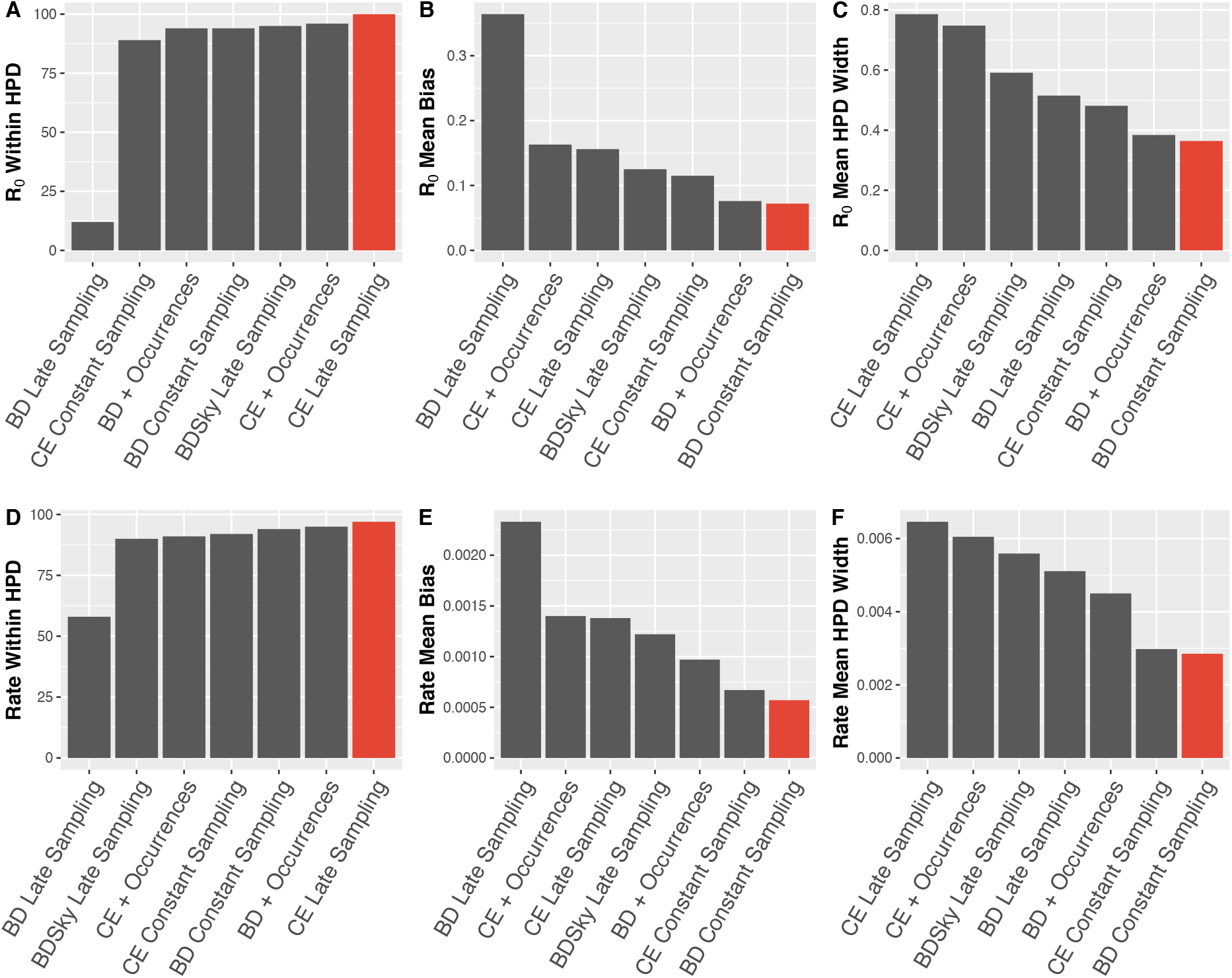
Bar ordering varies across plots to reflect preferential performance in each statistic such that those in red are most preferential. A) The number of simulations (out of 100) for which HPDs for R_0_ captured 2, the value simulated under. B) Mean bias in R_0_ across simulation treatments. C) Mean HPD width in R_0_ across simulation treatments. D) The number of simulations (out of 100) for which HPDs for evolutionary rate captured 0.01, the value simulated under. E) Mean bias in Rate across simulation treatments. F) Mean HPD width in rate across simulation treatments.

In general, we observed that the birth-death model tended to overestimate *R_0_* while the coalescent exponential underestimated it for data sets with late sampling (Fig 2). Estimates of the evolutionary rate displayed an identical pattern to those of *R_0_*, with the coalescent exponential and the birth-death model being the most and least accurate respectively at the expense of precision. However, the evolutionary rate appeared overall robust to the choice of the tree prior, with the only treatment producing a less than 90% coverage being the birth-death model with late sampling. This is a valuable consideration for analyses of future outbreaks as considerable attention is initially devoted to estimating a reliable evolutionary rate for a given pathogen because this is key to phylodynamic inference (Duchene et al., 2020).

As expected, analyses of the data with constant sampling were accurate in a majority of cases, with 94 and 89 out of 100 simulations covering *R_0_* alongside 94 and 92 for the evolutionary rate under the birth-death and the coalescent exponential models, respectively. The true model is the birth-death, and as such it is expected to perform better than the coalescent. Estimates of *R_0_* including occurrence data were similar in accuracy to those with complete sampling. A total of 94 analyses correctly estimated this parameter under the birth-death model, and 96 analyses included the true value for the coalescent exponential. Evolutionary rate estimates with occurrence data were similar, with 95 accurate estimates using the birth-death model and 91 using the coalescent exponential (Table 1, Fig2a,d). These results are attributable to the fact that the birth-death model treats sampling times as data, whereas the coalescent exponential model conditions on the number of samples and their ages (Boskova et al., 2018; Stadler et al., 2015). In the birth-death model, occurrence data improve accuracy when inferring *R_0_* and are also informative about the age of the tree height under this tree prior, which can also improve the accuracy of the evolutionary rate relative to the coalescent exponential model. But these estimates are unlikely to be as accurate as those with complete sequence data because they include less information.

The coalescent exponential model appears to be more robust to the sampling treatment, with greater accuracy than the birth-death model across late sampling and occurrence treatments. Our simulations suggest that this comes at the expense of less precise estimates than those from the birth-death model (Table 1). In turn, birth-death and birth-death skyline models tend to produce more precise estimates with less bias (Table 1, Fig 2). Together these results suggest that in a genomic-reporting scenario, the coalescent exponential is suitable when sampling proportion is assumed to be low, when the sampling process is otherwise poorly understood, or when no reliable occurrence data are available. However, when increased precision is desirable and occurrence data are available, birth-death tree priors may provide the sharper estimates with comparable accuracy. The choice of tree prior could be optimised depending on prioritisation of precision and bias based on the ordering of bars in Figure 2.

### Empirical case study: SARS-CoV-2 from the northwest USA

Mirroring trends in our simulated data sets, the coalescent exponential returned consistent estimates of *R_0_* across treatments which were generally lower than those inferred by the birth-death tree prior (Fig 3a, Table 2). Coalescent exponential treatments again produced wider HPD intervals than birth-death treatments, with the exception of late sampling which was highly uncertain under the birth-death, as expected from simulations. Uncertainty in posterior *R_0_* does not appear to change when substituting sequenced data for occurrence data (Fig 3A), indicating that late samples are highly informative while occurrence data contribute relatively little additional information to coalescent analyses. Moreover, we observed a near perfect match between estimates from analyses with only late sampling and those that included occurrences. This pattern can be explained because occurrence data have no influence on marginal posterior estimates under the coalescent. By contrast, our simulations show small differences in performance between coalescent analyses with late sampling and those with occurrence data, which we attribute to noise in the simulation study.

**Table 2.** Posterior estimates of *R_0_* and *p* using the birth-death for the SARS-CoV-2 empirical dataset. Rows correspond to the 12 treatments.

| Sampling Treatment | Mean $R_0$ | 95% HPD |
| --- | --- | --- |
| BD Constant Sampling | 1.96 | (1.85, 2.07) |
| BD Post 2020.18 | 2.44 | (2.31, 2.58) |
| BD Post 2020.18 + Occurrences | 1.97 | (1.87, 2.08) |
| BD Post 2020.2 | 3.53 | (3.24, 3.82) |
| BD Post 2020.2 + Occurrences | 1.96 | (1.83, 2.09) |
| BDSky Post 2020.18 | 1.57 | (1.43, 1.71) |
| BDSky Post 2020.2 | 1.48 | (1.35, 1.63) |
| CE Constant Sampling | 1.52 | (1.4, 1.65) |
| CE Post 2020.18 | 1.51 | (1.39, 1.64) |
| CE Post 2020.18 + Occurrences | 1.50 | (1.38, 1.62) |
| CE Post 2020.2 | 1.43 | (1.3, 1.58) |
| CE Post 2020.2 + Occurrences | 1.43 | (1.29, 1.57) |

**Fig 3.**
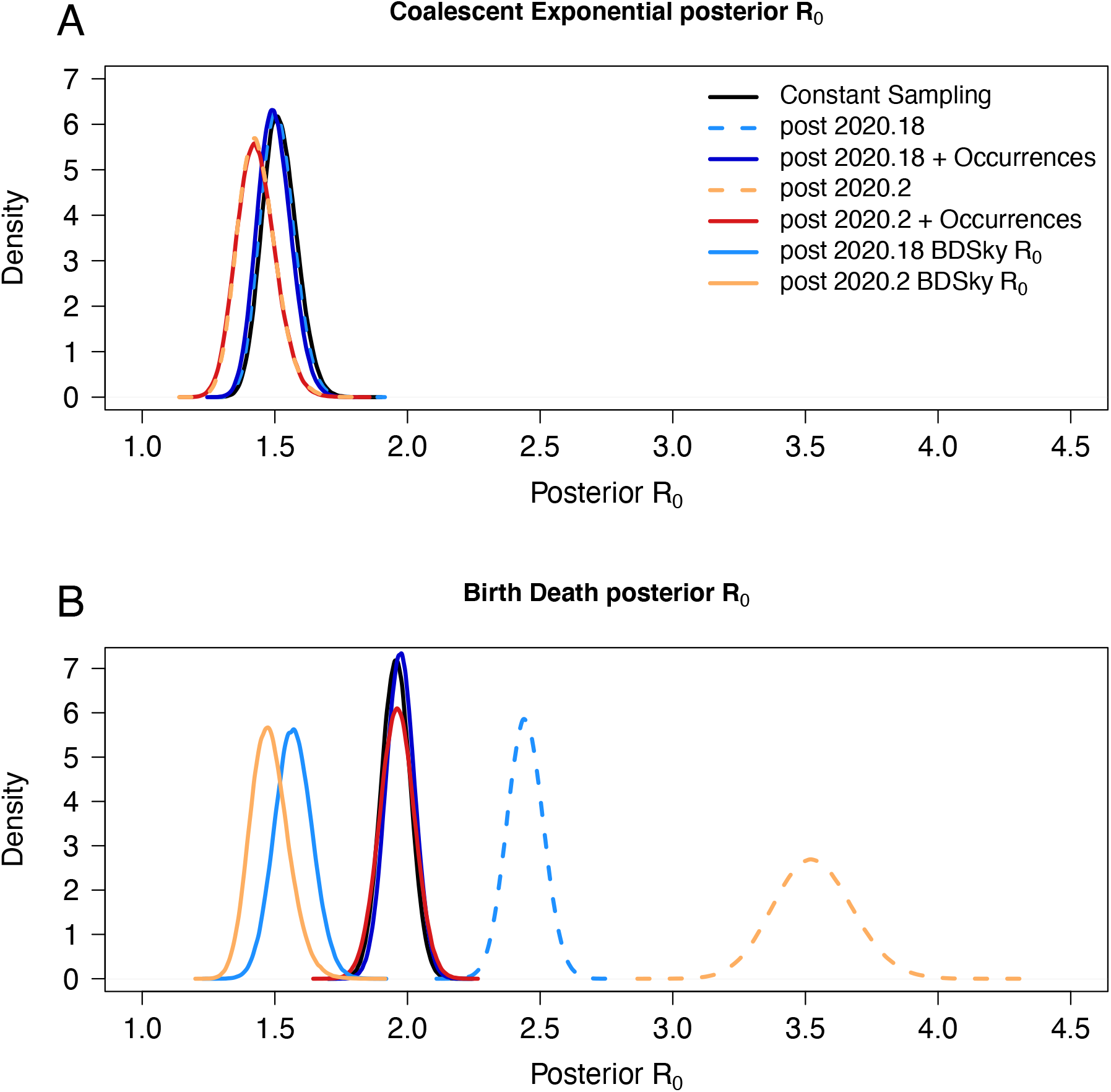
Posterior estimates of R_0_ for SARS-CoV-2 genome data. Constant sampling refers to using all 821 genomes in the empirical dataset. Post 2020.18 refers to only including sequences from 2020-03-04 and afterwards. Post 2020.2 refers to the same from 2020-03-14 and afterwards. A) Posterior densities of the basic reproductive number, *R_0_* under the coalescent exponential. B) Posterior densities for estimates of the basic reproductive number, *R_0_* under the birth death. In B, birth-death and birth-death skyline posteriors for *R_0_* and post cut-off sampling proportions are overlapping.

The results of the birth-death analyses recapitulate our observation from simulations that later sampling inflates estimates of *R_0_*, and that occurrence data rectify this (Figure 3B table 2). Complete sampling gave a mean *R_0_* of 1.96 (95% HPD: 1.85, 2.07) and late sampling with occurrence data estimated mean *R_0_* of 1.95 and 2.00 (95% HPDs: 1.8, 2.11 and 1.9, 2.12 for post - 2020.18 and 2020.2 respectively). These estimates are slightly lower than those from earlier work to estimate *R_0_* in the Washington state epidemic (Vaughan et al., 2020). This discrepancy may be due to the former being conducted earlier when the virus may have been spreading more rapidly. Late sampling alone inferred a mean *R_0_* of 2.44 and 3.53 for post 2020.18 and 2020.2 (2.31, 2.58 and 3.24, 3.82 95% HPDs respectively). The way in which the latest sampling data set inferred the highest values of *R_0_* further suggests that upward bias increases with lateness in sampling.

In both late sampling treatments, the birth-death skyline posterior distributions of *R_0_* were lower than their equivalents under the standard birth-death model, with later sampling corresponding to lower estimates (Fig 3). This is consistent with the simulated data (Fig S2), and suggests that including occurrence data is a preferential strategy to rectify posterior *R_0_* estimates amid late genome sequence sampling. Furthermore, the entropy of each birth-death based posterior *R_0_* distribution, a measure of uncertainty, is comparable at 3.68-3.78 as calculated with the mlf R package (Peterson, 2018). This further suggests that the topological uncertainty induced by occurrence data does not considerably increase uncertainty in posterior *R_0_* (Fig 3).

## Discussion

### Occurrence data in empirical phylodynamic studies

Occurrence data represent an extreme case of when genome coverage in samples is poor. Herein we show that low-coverage samples can be useful in phylodynamics so long as the sequences analysed are accurate. An outstanding task is to characterise an upper-bound on the relative proportion of occurrence to genomic samples from which genomic samples can still inform tree topology for epidemiological dynamics. To this end, we caution against over-inflating occurrence among genomic data sets without comparing to results obtained with genomic samples alone.

Our simulations and empirical data analyses reveal that occurrence data are a rich source of information for birth-death tree priors that can dramatically improve the accuracy and precision in estimates of epidemiological parameters. A key consideration is that occurrences should represent confirmed cases that would have been sequenced if sequencing effort had been constant, and which are known to belong to a particular outbreak, such as via contact tracing. Combining occurrence and sequence data can be particularly useful in situations where it is unknown if sequence sampling has been constant over time or where there exist several confirmed cases but a smaller number of sequences. This is valuable amid recently emerging outbreaks where combining both sources of data can provide sharper and more timely insight into the recent evolution of the pathogen in question.

## Acknowledgements

We are grateful to Trevor Bedford, authors and groups, originating laboratories, and submitting laboratories of sequences downloaded from GISAID. We provide a full acknowledgement table in the supplementary information. SD and LAF were supported by a Discovery Early Career Fellowship from the Australian Research Council (DE190100805), awarded to SD. ECH is supported by an Australian Research Council Laureate Fellowship (FL170100022).

## Authors’ contributions

All authors contributed to the design of experiments and writing of the manuscript. LF conducted analyses of empirical data and lead writing of the manuscript. FG contributed initial datasets, writing, and guidance with figures. TV contributed to writing the manuscript and mathematical concepts. EH contributed to writing of the manuscript and original ideas. SD conceived of fundamental concepts in the manuscript, conducted simulations, and contributed to writing.

## Data availability

Input files to generate trees in MASTER and to analyse sequence data in BEAST according to the birth-death skyline, birth-death, and the coalescent exponential tree priors, and accession numbers for empirical SARS-CoV-2 virus data. Available at: github.com/sebastianduchene/birth-death-sampling. Accession numbers for empirical SARS-CoV-2 virus data and the GISAID acknowledgements table are available as supplementary data online.

## Supplementary material

**Table S1.** Results of the simulation study with *R_0_*=1.5, evolutionary rate of 0.01 subs/site/year, and late sampling starting at 0.9 years of a total time of 1 year. The rows correspond to the seven treatments. The first two columns denote the number of simulations (out of 100) where the value used to generate the data was contained within the 95% highest posterior density (HPD). The last two columns are a measure of precision of the estimates calculated as the estimated mean estimate of *R_0_* and the evolutionary rate divided by the 95% HPD width, such that large values imply low precision. Here we report the mean value over 100 simulations.

| | $R_0$ within 95% HPD | Evol. rate within 95% HPD | Mean $R_0$ / HPD width | Mean evol. rate / HPD width |
| --- | --- | --- | --- | --- |
| Late sampling BD const. | 0 | 37 | 0.20 | 0.39 |
| Late sampling BD skyline | 92 | 93 | 0.25 | 0.54 |
| Late sampling Coal. exp. | 80 | 87 | 0.27 | 0.59 |
| Constant sampling BD const. | 97 | 94 | 0.16 | 0.23 |
| Constant sampling Coal. exp. | 96 | 92 | 0.16 | 0.23 |
| Birth Death + Occurrences. | 92 | 86 | 0.16 | 0.35 |
| Coalescent Exponential + Occurrences | 66 | 69 | 0.23 | 0.43 |

**Fig S1.**
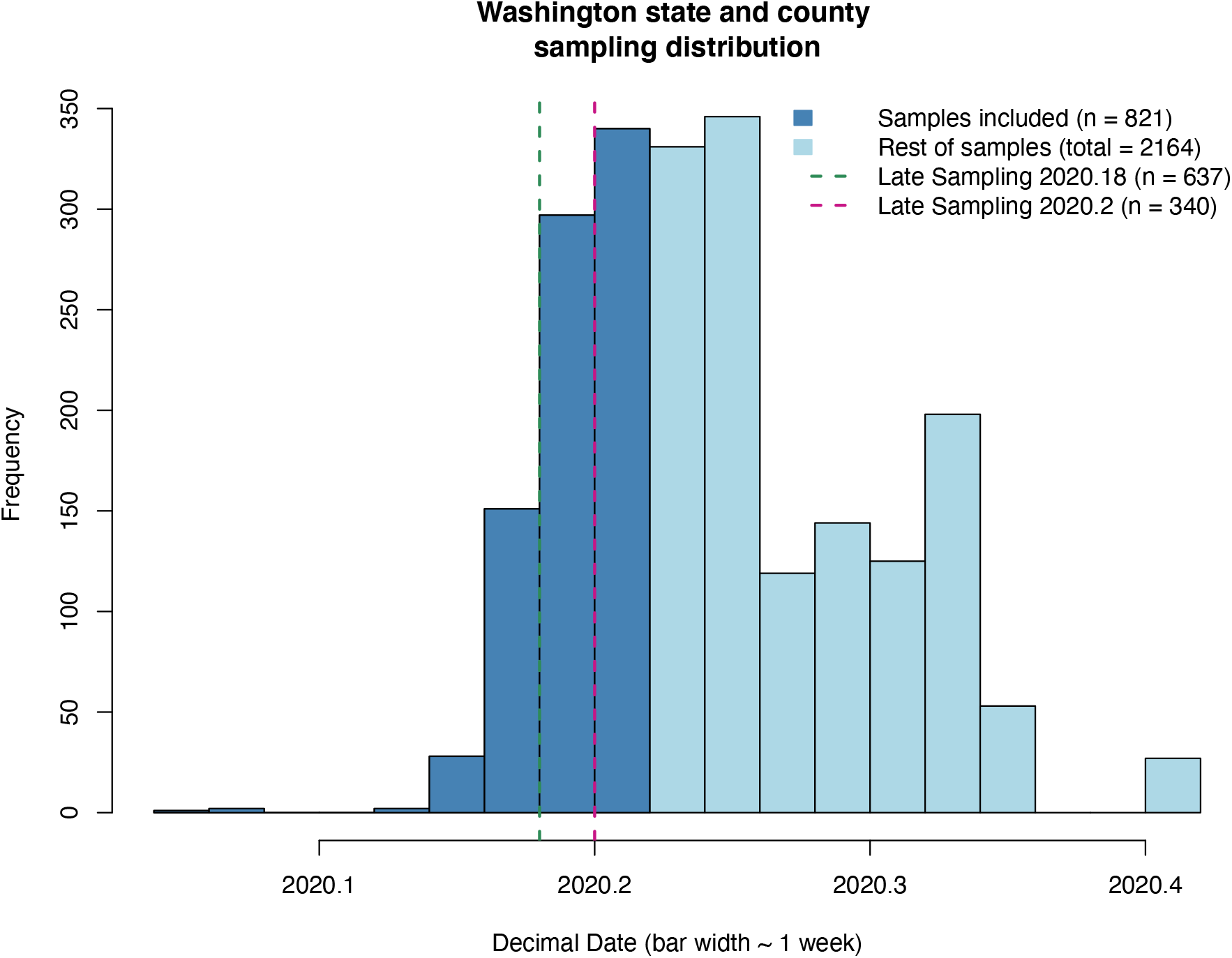
The temporal distribution of SARS-CoV-2 samples taken from Washington State and Washington County, Oregon, during the COVID-19 pandemic downloaded from GISAID. Colouring represents the subset of these data that we analysed and vertical lines show our two cut-offs for late sampling.

**Fig S2.**
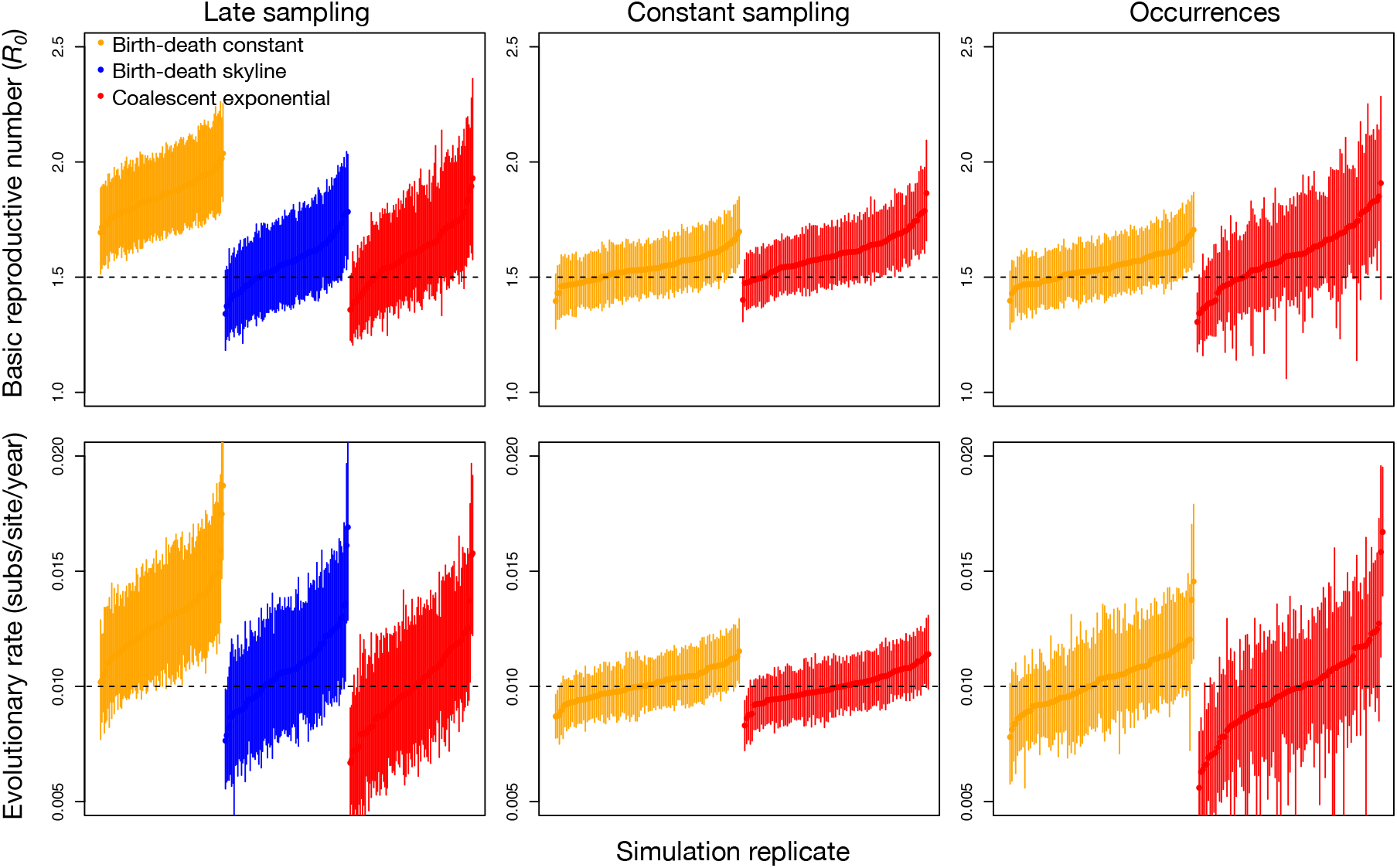
Posterior densities for estimates of the basic reproductive number, *R_0_*, and the evolutionary rate for 100 simulations with true *R_0_* of 1.5 and an evolutionary rate of 0.01 subs/site/year. The bars represent the 95% highest posterior density (HPD) and the points are the mean. Estimates are ordered from lowest to highest mean. We analysed the data by sampling late in the outbreak only (i.e. after 0.75 of the tree height), with a constant sampling effort (with all samples sequenced), and by including occurrence data. The colours represent four different tree priors; red for the coalescent exponential, blue for the birth-death skyline, and orange for the birth-death with constant sampling. For the data with sampling late in the outbreak only we use the birth-death skyline tree prior with constant *R_0_* and two intervals for the sampling rate, *ψ*, before time 0.75. This tree priori not applicable to analyses with complete sampling or with occurrence data where sampling is constant. The dashed horizontal lines correspond to the true parameter value used to generate the data.

