## Supplementary material for "Infectious disease phylodynamics with occurrence data"

We gratefully acknowledge the following Authors from the Originating laboratories responsible for obtaining the specimens, as well as the Submitting laboratories where the genome data were generated and shared via GISAID, on which this research is based.

All Submitters of data may be contacted directly via www.gisaid.org

| Accession ID | Originating Laboratory | Submitting Laboratory | Authors |
|---|---|---|---|
| EPI_ISL_412970 | Washington State Department of Health | Seattle Flu Study | Helen Chu, Michael Boeckh, Janet Englund, Michael Famulare, Barry Lutz, Deborah Nickerson, Mark Rieder, Lea Starita, Matthew Thompson, Jay Shendure, and Trevor Bedford |
| EPI_ISL_413025 | Harborview Medical Center | UW Virology Lab | Pavitra Roychoudhury, Arun Nalla, Hong Xie, Keith Jerome, Alexander Greninger |
| EPI_ISL_413455 | Washington State Public Health Lab | University of Washington Virology Lab | Pavitra Roychoudhury, Arun Nalla, Hong Xie, Keith Jerome, Alexander Greninger |
| EPI_ISL_413456 | Seattle Flu Study, University of Washington Medical Center | Seattle Flu Study, University of Washington Medical Center | Chu et al |
| EPI_ISL_413457, EPI_ISL_413458 | Washington State Public Health Lab | UW Virology Lab | Pavitra Roychoudhury, Arun Nalla, Hong Xie, Keith Jerome, Alexander Greninger |
| EPI_ISL_413486 | Valley Medical Center | University of Washington Virology Lab | Pavitra Roychoudhury, Arun Nalla, Hong Xie, Keith Jerome, Alexander Greninger |
| EPI_ISL_413487 | Harborview Medical Center | University of Washington Virology Lab | Pavitra Roychoudhury, Arun Nalla, Hong Xie, Keith Jerome, Alexander Greninger |
| EPI_ISL_413560 | Seattle Flu Study | Seattle Flu Study | Chu et al |
| EPI_ISL_413562, EPI_ISL_413563 | UW Virology Lab | UW Virology Lab | Pavitra Roychoudhury, Hong Xie, Keith Jerome, Alexander Greninger |
| EPI_ISL_413601 | UW Virology Lab | UW Virology Lab | Pavitra Roychoudhury, Hong Xie, Keith Jerome, Alexander Greninger |
| EPI_ISL_413649, EPI_ISL_413650, EPI_ISL_413651, EPI_ISL_413652, EPI_ISL_413653, EPI_ISL_414363, EPI_ISL_414364, EPI_ISL_414365, EPI_ISL_414366, EPI_ISL_414367, EPI_ISL_414368, EPI_ISL_414369, EPI_ISL_414591, EPI_ISL_414592, EPI_ISL_414593, EPI_ISL_414594, EPI_ISL_414595, EPI_ISL_414596, EPI_ISL_414597, EPI_ISL_414616, EPI_ISL_414621, EPI_ISL_414622, EPI_ISL_415591, EPI_ISL_415592, EPI_ISL_415593, EPI_ISL_415594, EPI_ISL_415595, EPI_ISL_415596, EPI_ISL_415597, EPI_ISL_415598, EPI_ISL_415599, EPI_ISL_415601, EPI_ISL_415602, EPI_ISL_415603, EPI_ISL_415604, EPI_ISL_415605, EPI_ISL_415606, EPI_ISL_415607, EPI_ISL_415608, EPI_ISL_415609, EPI_ISL_415610, EPI_ISL_415611, EPI_ISL_415612, EPI_ISL_415613, EPI_ISL_415614, EPI_ISL_415615, EPI_ISL_415616, EPI_ISL_415617, EPI_ISL_415618, EPI_ISL_415619, EPI_ISL_415620, EPI_ISL_415621, EPI_ISL_415622, EPI_ISL_415623, EPI_ISL_415624, EPI_ISL_415625, EPI_ISL_415626, EPI_ISL_415627, EPI_ISL_416433, EPI_ISL_416434, EPI_ISL_416435, EPI_ISL_416436, EPI_ISL_416437, EPI_ISL_416438, EPI_ISL_416439, EPI_ISL_416440, EPI_ISL_416441, EPI_ISL_416442, EPI_ISL_416443, EPI_ISL_416444, EPI_ISL_416445, EPI_ISL_416446, EPI_ISL_416447, EPI_ISL_416448, EPI_ISL_416449, EPI_ISL_416450, EPI_ISL_416451, EPI_ISL_416452, EPI_ISL_416453, EPI_ISL_416454, EPI_ISL_416455, EPI_ISL_416456 | | | |
| see above | UW Virology Lab | UW Virology Lab | Pavitra Roychoudhury, Hong Xie, Keith Jerome, Alexander Greninger |
| EPI_ISL_416459, EPI_ISL_416460, EPI_ISL_416461, EPI_ISL_416462, EPI_ISL_416463, EPI_ISL_416464, EPI_ISL_416465, EPI_ISL_416466 | Seattle Flu Study | Seattle Flu Study | Chu et al |
| EPI_ISL_416635, EPI_ISL_416636, EPI_ISL_416637, EPI_ISL_416638, EPI_ISL_416639, EPI_ISL_416640, EPI_ISL_416641, EPI_ISL_416642, EPI_ISL_416643, EPI_ISL_416644, EPI_ISL_416645, EPI_ISL_416646, EPI_ISL_416647, EPI_ISL_416648, EPI_ISL_416649, EPI_ISL_416652, EPI_ISL_416653, EPI_ISL_416654, EPI_ISL_416655, EPI_ISL_416656, EPI_ISL_416657, EPI_ISL_416658, EPI_ISL_416659, EPI_ISL_416661, EPI_ISL_416662, EPI_ISL_416663, EPI_ISL_416664, EPI_ISL_416665, EPI_ISL_416666, EPI_ISL_416667, EPI_ISL_416668, EPI_ISL_416669, EPI_ISL_416670, EPI_ISL_416671, EPI_ISL_416677, EPI_ISL_416678, EPI_ISL_416679, EPI_ISL_416680, EPI_ISL_416681, EPI_ISL_416683, EPI_ISL_416684, EPI_ISL_416685, EPI_ISL_416686, EPI_ISL_416687, EPI_ISL_416688, EPI_ISL_416689, EPI_ISL_416690, EPI_ISL_416691, EPI_ISL_416692, EPI_ISL_416693, EPI_ISL_416694, EPI_ISL_416695, EPI_ISL_416698, EPI_ISL_416699, EPI_ISL_416700, EPI_ISL_416701, EPI_ISL_416702, EPI_ISL_416703, EPI_ISL_416704, EPI_ISL_416705, EPI_ISL_416706, EPI_ISL_416707, EPI_ISL_416708, EPI_ISL_416709, EPI_ISL_416710, EPI_ISL_416711, EPI_ISL_416712, EPI_ISL_416713, EPI_ISL_416714, EPI_ISL_416715, EPI_ISL_416716, EPI_ISL_416717, EPI_ISL_416718, EPI_ISL_416719, EPI_ISL_416720, EPI_ISL_416721, EPI_ISL_416722, EPI_ISL_416723, EPI_ISL_416724, EPI_ISL_416725, EPI_ISL_416726, EPI_ISL_416727, EPI_ISL_416728, EPI_ISL_416729 | | | |
| see above | UW Virology Lab | UW Virology Lab | Pavitra Roychoudhury, Hong Xie, Keith Jerome, Alexander Greninger |
| EPI_ISL_417065, EPI_ISL_417066, EPI_ISL_417067, EPI_ISL_417068, EPI_ISL_417069, EPI_ISL_417070, EPI_ISL_417071, EPI_ISL_417072, EPI_ISL_417073, EPI_ISL_417074, EPI_ISL_417075, EPI_ISL_417076, EPI_ISL_417077, EPI_ISL_417078, EPI_ISL_417079, EPI_ISL_417080, EPI_ISL_417081, EPI_ISL_417082, EPI_ISL_417083, EPI_ISL_417084, EPI_ISL_417085, EPI_ISL_417086, EPI_ISL_417087, EPI_ISL_417088, EPI_ISL_417089, EPI_ISL_417090, EPI_ISL_417091, EPI_ISL_417092, EPI_ISL_417093, EPI_ISL_417094, EPI_ISL_417095, EPI_ISL_417096, EPI_ISL_417097, EPI_ISL_417098, EPI_ISL_417099, EPI_ISL_417100, EPI_ISL_417101, EPI_ISL_417102, EPI_ISL_417103, EPI_ISL_417104, EPI_ISL_417105, EPI_ISL_417106, EPI_ISL_417107, EPI_ISL_417108, EPI_ISL_417109, EPI_ISL_417110, EPI_ISL_417111, EPI_ISL_417112, EPI_ISL_417113, EPI_ISL_417114, EPI_ISL_417115, EPI_ISL_417116, EPI_ISL_417117, EPI_ISL_417118, EPI_ISL_417119, EPI_ISL_417120, EPI_ISL_417121, EPI_ISL_417122, EPI_ISL_417123, EPI_ISL_417124, EPI_ISL_417125, EPI_ISL_417126, EPI_ISL_417127, EPI_ISL_417128, EPI_ISL_417129, EPI_ISL_417130, EPI_ISL_417131, EPI_ISL_417132, EPI_ISL_417133, EPI_ISL_417134, EPI_ISL_417135, EPI_ISL_417136, EPI_ISL_417137, EPI_ISL_417138, EPI_ISL_417139, EPI_ISL_417140, EPI_ISL_417141, EPI_ISL_417142, EPI_ISL_417143, EPI_ISL_417144, EPI_ISL_417145, EPI_ISL_417146, EPI_ISL_417147, EPI_ISL_417148, EPI_ISL_417149, EPI_ISL_417150, EPI_ISL_417151, EPI_ISL_417152, EPI_ISL_417153, EPI_ISL_417154, EPI_ISL_417155, EPI_ISL_417156, EPI_ISL_417157, EPI_ISL_417158, EPI_ISL_417159, EPI_ISL_417160, EPI_ISL_417161, EPI_ISL_417162 | | | |
| see above | Washington State Department of Health | Seattle Flu Study | Chu etl al |
| EPI_ISL_417163, EPI_ISL_417164, EPI_ISL_417165 | Seattle Flu Study | Seattle Flu Study | Chu etl al |
| EPI_ISL_417166, EPI_ISL_417167, EPI_ISL_417168, EPI_ISL_417169, EPI_ISL_417170, EPI_ISL_417171, EPI_ISL_417172, EPI_ISL_417173, EPI_ISL_417174, EPI_ISL_417175 | Washington State Department of Health | Seattle Flu Study | Chu etl al |
| EPI_ISL_417341, EPI_ISL_417342, EPI_ISL_417343, EPI_ISL_417344, EPI_ISL_417345, EPI_ISL_417346, EPI_ISL_417347, EPI_ISL_417348, EPI_ISL_417349, EPI_ISL_417350, EPI_ISL_417351, EPI_ISL_417352, EPI_ISL_417353, EPI_ISL_417354, EPI_ISL_417355, EPI_ISL_417356, EPI_ISL_417357, EPI_ISL_417358, EPI_ISL_417359, EPI_ISL_417360, EPI_ISL_417361, EPI_ISL_417363, EPI_ISL_417364, EPI_ISL_417365, EPI_ISL_417366, EPI_ISL_417367, EPI_ISL_417369, EPI_ISL_417370, EPI_ISL_417371, EPI_ISL_417374, EPI_ISL_417375, EPI_ISL_417376, EPI_ISL_417377, EPI_ISL_417378, EPI_ISL_417379, EPI_ISL_417380, EPI_ISL_417381, EPI_ISL_417382, EPI_ISL_417448, EPI_ISL_417449, EPI_ISL_417450, EPI_ISL_417451, EPI_ISL_417453, EPI_ISL_417454, EPI_ISL_417455, EPI_ISL_417456, EPI_ISL_418029, EPI_ISL_418030, EPI_ISL_418031, EPI_ISL_418032, EPI_ISL_418033, EPI_ISL_418034, EPI_ISL_418037, EPI_ISL_418038, EPI_ISL_418040, EPI_ISL_418046, EPI_ISL_418047, EPI_ISL_418048, EPI_ISL_418050, EPI_ISL_418052, EPI_ISL_418053, EPI_ISL_418054, EPI_ISL_418055, EPI_ISL_418062, EPI_ISL_418063, EPI_ISL_418064, EPI_ISL_418065, EPI_ISL_418066, EPI_ISL_418067, EPI_ISL_418068, EPI_ISL_418069, EPI_ISL_418070, EPI_ISL_418071, EPI_ISL_418072, EPI_ISL_418073, EPI_ISL_418074, EPI_ISL_418075, EPI_ISL_418076, EPI_ISL_418077, EPI_ISL_418078, EPI_ISL_418079, EPI_ISL_418080, EPI_ISL_418081, EPI_ISL_418082 | | | |
| see above | UW Virology Lab | UW Virology Lab | Pavitra Roychoudhury, Hong Xie, Keith Jerome, Alexander Greninger |
| EPI_ISL_418771, EPI_ISL_418772, EPI_ISL_418773, EPI_ISL_418774, EPI_ISL_418775, EPI_ISL_418776, EPI_ISL_418777 | WA State Department of Health | Pathogen Discovery, Respiratory Viruses Branch, Division of Viral Diseases, Centers for Disease Control and Prevention | Jing Zhang, Ying Tao, Clinton R. Paden, Krista Queen, Anna Uehara, Yan Li, Haibin Wang, Jesica Jacobs, Denny Russell, Brian Hiatt, Jessica Gant, Suxiang Tong |
| EPI_ISL_418778, EPI_ISL_418779, EPI_ISL_418780, EPI_ISL_418781, EPI_ISL_418782, EPI_ISL_418783, EPI_ISL_418784, EPI_ISL_418785, EPI_ISL_418786, EPI_ISL_418787, EPI_ISL_418788, EPI_ISL_418789, EPI_ISL_418790, EPI_ISL_418791 | | | |
| see above | WA State Department of Health | Pathogen Discovery, Respiratory Viruses Branch, Division of Viral Diseases, Centers for Disease Control and Prevention | Ying Tao, Jing Zhang, Clinton R. Paden, Krista Queen, Anna Uehara, Yan Li, Haibin Wang, Jesica Jacobs, Denny Russell, Brian Hiatt, Jessica Gant, Suxiang Tong |
| EPI_ISL_418866, EPI_ISL_418867, EPI_ISL_418868, EPI_ISL_418869, EPI_ISL_418870, EPI_ISL_418872, EPI_ISL_418873, EPI_ISL_418874, EPI_ISL_418875, EPI_ISL_418876, EPI_ISL_418877, EPI_ISL_418878, EPI_ISL_418879, EPI_ISL_418880, EPI_ISL_418881, EPI_ISL_418882, EPI_ISL_418883, EPI_ISL_418884, EPI_ISL_418885, EPI_ISL_418886, EPI_ISL_418887, EPI_ISL_418888, EPI_ISL_418889, EPI_ISL_418890, EPI_ISL_418891, EPI_ISL_418892, EPI_ISL_418897, EPI_ISL_418899, EPI_ISL_418900, EPI_ISL_418903, EPI_ISL_418904, EPI_ISL_418905, EPI_ISL_418906, EPI_ISL_418907, EPI_ISL_418908, EPI_ISL_418909, EPI_ISL_418910, EPI_ISL_418911, EPI_ISL_418912, EPI_ISL_418913, EPI_ISL_418914, EPI_ISL_418922, EPI_ISL_418923, EPI_ISL_418924, EPI_ISL_418926, EPI_ISL_418927, EPI_ISL_418928, EPI_ISL_418929, EPI_ISL_418930, EPI_ISL_418931, EPI_ISL_418932, EPI_ISL_418933, EPI_ISL_418934, EPI_ISL_418935, EPI_ISL_418936, EPI_ISL_418937, EPI_ISL_418938, EPI_ISL_418939, EPI_ISL_418940, EPI_ISL_418942, EPI_ISL_418943, EPI_ISL_418944, EPI_ISL_418945, EPI_ISL_418948, EPI_ISL_418949, EPI_ISL_418950, EPI_ISL_418951, EPI_ISL_418952, EPI_ISL_418953, EPI_ISL_418954, EPI_ISL_418955 | | | |
| see above | UW Virology Lab | UW Virology Lab | Pavitra Roychoudhury, Hong Xie, Keith Jerome, Alexander Greninger |
| EPI_ISL_419555 | WA State Department of Health | Pathogen Discovery, Respiratory Viruses Branch, Division of Viral Diseases, Centers for Disease Control and Prevention | Ying Tao, Jing Zhang, Krista Queen, Anna Uehara, Clinton R. Paden, Yan Li, Haibin Wang, Jasmine Padilla, Justin Lee, Suxiang Tong |
| EPI_ISL_422963, EPI_ISL_422964, EPI_ISL_422965, EPI_ISL_422966, EPI_ISL_422967, EPI_ISL_422969, EPI_ISL_422970, EPI_ISL_422971, EPI_ISL_422972, EPI_ISL_422973, EPI_ISL_422974, EPI_ISL_422975, EPI_ISL_422976, EPI_ISL_422977, EPI_ISL_422981, EPI_ISL_422984, EPI_ISL_422985, EPI_ISL_422986, EPI_ISL_422987, EPI_ISL_422988, EPI_ISL_422989, EPI_ISL_422990, EPI_ISL_422991, EPI_ISL_422992, EPI_ISL_422993, EPI_ISL_422994, EPI_ISL_422995, EPI_ISL_422996, EPI_ISL_422997, EPI_ISL_422998, EPI_ISL_422999, EPI_ISL_423000, EPI_ISL_423001, EPI_ISL_423002, EPI_ISL_423003, EPI_ISL_423004, EPI_ISL_423005, EPI_ISL_423006, EPI_ISL_423007, EPI_ISL_423008, EPI_ISL_423009, EPI_ISL_423010, EPI_ISL_423011, EPI_ISL_423012, EPI_ISL_423013, EPI_ISL_423014, EPI_ISL_423015, EPI_ISL_423016, EPI_ISL_423017, EPI_ISL_423018, EPI_ISL_423019, EPI_ISL_423020, EPI_ISL_423021, EPI_ISL_423022, EPI_ISL_423024, EPI_ISL_423025, EPI_ISL_423026, EPI_ISL_423027, EPI_ISL_423028, EPI_ISL_423029, EPI_ISL_423030, EPI_ISL_423031, EPI_ISL_423032, EPI_ISL_423033, EPI_ISL_424166, EPI_ISL_424167, EPI_ISL_424168, EPI_ISL_424169, EPI_ISL_424170, EPI_ISL_424171, EPI_ISL_424172, EPI_ISL_424175, EPI_ISL_424176, EPI_ISL_424177, EPI_ISL_424178, EPI_ISL_424179, EPI_ISL_424180, EPI_ISL_424182, EPI_ISL_424183, EPI_ISL_424184, EPI_ISL_424186, EPI_ISL_424187, EPI_ISL_424188, EPI_ISL_424190, EPI_ISL_424191, EPI_ISL_424192, EPI_ISL_424193, EPI_ISL_424194, EPI_ISL_424195, EPI_ISL_424196, EPI_ISL_424197, EPI_ISL_424198, EPI_ISL_424199, EPI_ISL_424200, EPI_ISL_424201, EPI_ISL_424202, EPI_ISL_424203, EPI_ISL_424204, EPI_ISL_424205, EPI_ISL_424206, EPI_ISL_424207, EPI_ISL_424208, EPI_ISL_424209, EPI_ISL_424210, EPI_ISL_424211, EPI_ISL_424212, EPI_ISL_424213, EPI_ISL_424214, EPI_ISL_424215, EPI_ISL_424216, EPI_ISL_424217, EPI_ISL_424218, EPI_ISL_424219, EPI_ISL_424220, EPI_ISL_424221, EPI_ISL_424222, EPI_ISL_424223, EPI_ISL_424224, EPI_ISL_424225, EPI_ISL_424226, EPI_ISL_424227, EPI_ISL_424228, EPI_ISL_424229, EPI_ISL_424230, EPI_ISL_424231, EPI_ISL_424232, EPI_ISL_424233, EPI_ISL_424234, EPI_ISL_424235, EPI_ISL_424236, EPI_ISL_424237, EPI_ISL_424238, EPI_ISL_424239, EPI_ISL_424240, EPI_ISL_424241, EPI_ISL_424242, EPI_ISL_424243, EPI_ISL_424244, EPI_ISL_424245, EPI_ISL_424246, EPI_ISL_424247, EPI_ISL_424250, EPI_ISL_424251, EPI_ISL_424252, EPI_ISL_424253, EPI_ISL_424254, EPI_ISL_424256, EPI_ISL_424257, EPI_ISL_424258, EPI_ISL_424259, EPI_ISL_424260, EPI_ISL_424261, EPI_ISL_424262, EPI_ISL_424263, EPI_ISL_424264, EPI_ISL_424269, EPI_ISL_424270, EPI_ISL_424271, EPI_ISL_424272, EPI_ISL_424273, EPI_ISL_424274, EPI_ISL_424275, EPI_ISL_424276, EPI_ISL_424277, EPI_ISL_424278, EPI_ISL_424279, EPI_ISL_424280, EPI_ISL_424281, EPI_ISL_424282, EPI_ISL_424283, EPI_ISL_424284, EPI_ISL_424287, EPI_ISL_424288, EPI_ISL_424289, EPI_ISL_424290, EPI_ISL_424291, EPI_ISL_424292, EPI_ISL_424293, EPI_ISL_424294, EPI_ISL_424295, EPI_ISL_424296, EPI_ISL_424297, EPI_ISL_424298, EPI_ISL_424299, EPI_ISL_424300, EPI_ISL_424301, EPI_ISL_424302, EPI_ISL_424304, EPI_ISL_424305, EPI_ISL_424306, EPI_ISL_424309, EPI_ISL_424310, EPI_ISL_424311, EPI_ISL_424312, EPI_ISL_424313, EPI_ISL_424314, EPI_ISL_424316, EPI_ISL_424317, EPI_ISL_424323, EPI_ISL_424324, EPI_ISL_424325, EPI_ISL_424326, EPI_ISL_424327, EPI_ISL_424328, EPI_ISL_424329, EPI_ISL_424330, EPI_ISL_424331, EPI_ISL_424332, EPI_ISL_424333, EPI_ISL_424334, EPI_ISL_424335, EPI_ISL_424336, EPI_ISL_424337, EPI_ISL_424338, EPI_ISL_424339, EPI_ISL_424340, EPI_ISL_424341, EPI_ISL_426079, EPI_ISL_426080, EPI_ISL_426081, EPI_ISL_426082, EPI_ISL_426083, EPI_ISL_426084, EPI_ISL_426085, EPI_ISL_426086, EPI_ISL_426087, EPI_ISL_426088, EPI_ISL_426089, EPI_ISL_426090, EPI_ISL_426091, EPI_ISL_426092, EPI_ISL_426093, EPI_ISL_426094, EPI_ISL_426095, EPI_ISL_426096, EPI_ISL_426097, EPI_ISL_426098, EPI_ISL_426099, EPI_ISL_426100, EPI_ISL_426101, EPI_ISL_426102, EPI_ISL_426103, EPI_ISL_426104, EPI_ISL_426105, EPI_ISL_426106, EPI_ISL_426107, EPI_ISL_426108, EPI_ISL_426109, EPI_ISL_426110, EPI_ISL_426111, EPI_ISL_426112, EPI_ISL_426113, EPI_ISL_426114, EPI_ISL_426115, EPI_ISL_426116, EPI_ISL_426117, EPI_ISL_426118, EPI_ISL_426119, EPI_ISL_426120, EPI_ISL_426121, EPI_ISL_426122, EPI_ISL_426123, EPI_ISL_426124, EPI_ISL_426125, EPI_ISL_426126, EPI_ISL_426127, EPI_ISL_426128, EPI_ISL_426129, EPI_ISL_426130, EPI_ISL_426131, EPI_ISL_426132, EPI_ISL_426133, EPI_ISL_426134, EPI_ISL_426135, EPI_ISL_426136, EPI_ISL_426137 | | | |
| see above | UW Virology Lab | UW Virology Lab | Pavitra Roychoudhury, Hong Xie, Keith Jerome, Alexander Greninger |
| EPI_ISL_426305 | Wadsworth Center, New York State Department.of Health | Wadsworth Center, New York State Department.of Health | Kirsten St. George, Daryl M. Lamson, Sara Griesemer, Jonathan Plitnick, Navjot Singh, Matthew D. Shudt, Erica Lasek-Nesselquist |
| EPI_ISL_426436 | WA State Department of Health | Pathogen Discovery, Respiratory Viruses Branch, Division of Viral Diseases, Centers for Disease Control and Prevention | Jing Zhang, Ying Tao, Clinton R. Paden, Krista Queen, Anna Uehara, Yan Li, Haibin Wang, Jesica Jacobs, Denny Russell, Brian Hiatt, Jessica Gant, Suxiang Tong |
| EPI_ISL_426437 | WA State Department of Health | Pathogen Discovery, Respiratory Viruses Branch, Division of Viral Diseases, Centers for Disease Control and Prevention | Ying Tao, Jing Zhang, Clinton R. Paden, Krista Queen, Anna Uehara, Yan Li, Haibin Wang, Jesica Jacobs, Denny Russell, Brian Hiatt, Jessica Gant, Suxiang Tong |
| EPI_ISL_426438, EPI_ISL_426439 | WA State Department of Health | Pathogen Discovery, Respiratory Viruses Branch, Division of Viral Diseases, Centers for Disease Control and Prevention | Jing Zhang, Ying Tao, Clinton R. Paden, Krista Queen, Anna Uehara, Yan Li, Haibin Wang, Jesica Jacobs, Denny Russell, Brian Hiatt, Jessica Gant, Suxiang Tong |
| EPI_ISL_426440, EPI_ISL_426441 | WA State Department of Health | Pathogen Discovery, Respiratory Viruses Branch, Division | Ying Tao, Jing Zhang, Clinton R. Paden, Krista Queen, Anna Uehara, Yan Li, Haibin Wang, Jesica Jacobs, Denny Russell, Brian Hiatt, Jessica Gant, Suxiang Tong |

| | | | |
|---|---|---|---|
| | | of Viral Diseases, Centers for Disease Control and Prevention | |
| EPI_ISL_426442 | WA State Department of Health | Pathogen Discovery, Respiratory Viruses Branch, Division of Viral Diseases, Centers for Disease Control and Prevention | Jing Zhang, Ying Tao, Clinton R. Paden, Krista Queen, Anna Uehara, Yan Li, Haibin Wang, Jesica Jacobs, Denny Russell, Brian Hiatt, Jessica Gant, Suxiang Tong |
| EPI_ISL_426443, EPI_ISL_426444 | WA State Department of Health | Pathogen Discovery, Respiratory Viruses Branch, Division of Viral Diseases, Centers for Disease Control and Prevention | Ying Tao, Jing Zhang, Clinton R. Paden, Krista Queen, Anna Uehara, Yan Li, Haibin Wang, Jesica Jacobs, Denny Russell, Brian Hiatt, Jessica Gant, Suxiang Tong |
| EPI_ISL_426445 | WA State Department of Health | Pathogen Discovery, Respiratory Viruses Branch, Division of Viral Diseases, Centers for Disease Control and Prevention | Jing Zhang, Ying Tao, Clinton R. Paden, Krista Queen, Anna Uehara, Yan Li, Haibin Wang, Jesica Jacobs, Denny Russell, Brian Hiatt, Jessica Gant, Suxiang Tong |
| EPI_ISL_426446, EPI_ISL_426447, EPI_ISL_426448, EPI_ISL_426449 | WA State Department of Health | Pathogen Discovery, Respiratory Viruses Branch, Division of Viral Diseases, Centers for Disease Control and Prevention | Ying Tao, Jing Zhang, Clinton R. Paden, Krista Queen, Anna Uehara, Yan Li, Haibin Wang, Jesica Jacobs, Denny Russell, Brian Hiatt, Jessica Gant, Suxiang Tong |
| EPI_ISL_426450 | WA State Department of Health | Pathogen Discovery, Respiratory Viruses Branch, Division of Viral Diseases, Centers for Disease Control and Prevention | Jing Zhang, Ying Tao, Clinton R. Paden, Krista Queen, Anna Uehara, Yan Li, Haibin Wang, Jesica Jacobs, Denny Russell, Brian Hiatt, Jessica Gant, Suxiang Tong |
| EPI_ISL_426451, EPI_ISL_426452, EPI_ISL_426453 | WA State Department of Health | Pathogen Discovery, Respiratory Viruses Branch, Division of Viral Diseases, Centers for Disease Control and Prevention | Ying Tao, Jing Zhang, Clinton R. Paden, Krista Queen, Anna Uehara, Yan Li, Haibin Wang, Jesica Jacobs, Denny Russell, Brian Hiatt, Jessica Gant, Suxiang Tong |
| EPI_ISL_427174, EPI_ISL_427175, EPI_ISL_427176, EPI_ISL_427177, EPI_ISL_427178, EPI_ISL_427179, EPI_ISL_427180, EPI_ISL_427181, EPI_ISL_427182, EPI_ISL_427183, EPI_ISL_427184, EPI_ISL_427185, EPI_ISL_427186, EPI_ISL_427187, EPI_ISL_427188, EPI_ISL_427189, EPI_ISL_427190, EPI_ISL_427191, EPI_ISL_427192, EPI_ISL_427193, EPI_ISL_427194, EPI_ISL_427195, EPI_ISL_427196, EPI_ISL_427197, EPI_ISL_427198, EPI_ISL_427199, EPI_ISL_427200, EPI_ISL_427201, EPI_ISL_427202, EPI_ISL_427203, EPI_ISL_427204, EPI_ISL_427205, EPI_ISL_427206, EPI_ISL_427207, EPI_ISL_427208, EPI_ISL_427209, EPI_ISL_427210, EPI_ISL_427211, EPI_ISL_427212, EPI_ISL_427213, EPI_ISL_427214, EPI_ISL_427215, EPI_ISL_427216, EPI_ISL_427217, EPI_ISL_427218, EPI_ISL_427219, EPI_ISL_427220, EPI_ISL_427221, EPI_ISL_427222, EPI_ISL_427223, EPI_ISL_427224, EPI_ISL_427225, EPI_ISL_427226, EPI_ISL_427227, EPI_ISL_427228, EPI_ISL_427229, EPI_ISL_427230, EPI_ISL_427231, EPI_ISL_427232, EPI_ISL_427233, EPI_ISL_427234, EPI_ISL_427235, EPI_ISL_427236, EPI_ISL_427237, EPI_ISL_427238, EPI_ISL_427239, EPI_ISL_427240, EPI_ISL_427241, EPI_ISL_427242, EPI_ISL_427243, EPI_ISL_427244, EPI_ISL_427245, EPI_ISL_427246, EPI_ISL_427247, EPI_ISL_427248, EPI_ISL_427249, EPI_ISL_427250, EPI_ISL_427251, EPI_ISL_427252, EPI_ISL_427253, EPI_ISL_427254, EPI_ISL_427255, EPI_ISL_427256, EPI_ISL_427257, EPI_ISL_427258, EPI_ISL_427259, EPI_ISL_427260, EPI_ISL_427261, EPI_ISL_427262, EPI_ISL_427263, EPI_ISL_427264, EPI_ISL_427265, EPI_ISL_427266, EPI_ISL_427267, EPI_ISL_427268, EPI_ISL_427269, EPI_ISL_427270 | | | |
| see above | UW Virology Lab | UW Virology Lab | Pavitra Roychoudhury, Hong Xie, Keith Jerome, Alexander Greninger |
| EPI_ISL_428289 | University of Wisconsin-Madison AIDS Vaccine Research Laboratories | University of Wisconsin-Madison AIDS Vaccine Research Laboratories | Gage Moreno, Katarina Braun, et al. AIDS Vaccine Research Laboratories |
| EPI_ISL_429597, EPI_ISL_429598, EPI_ISL_429600, EPI_ISL_429602, EPI_ISL_429603, EPI_ISL_429604, EPI_ISL_429605, EPI_ISL_429606, EPI_ISL_429607, EPI_ISL_429608, EPI_ISL_429609, EPI_ISL_429611, EPI_ISL_429612, EPI_ISL_429613, EPI_ISL_429614, EPI_ISL_429615, EPI_ISL_429616, EPI_ISL_429617, EPI_ISL_429618, EPI_ISL_429619, EPI_ISL_429620, EPI_ISL_429621, EPI_ISL_429622, EPI_ISL_429623, EPI_ISL_429624, EPI_ISL_429625, EPI_ISL_429626, EPI_ISL_429627, EPI_ISL_429629, EPI_ISL_429630, EPI_ISL_429631, EPI_ISL_429632, EPI_ISL_429633, EPI_ISL_429634, EPI_ISL_429635, EPI_ISL_429636, EPI_ISL_429637, EPI_ISL_429638, EPI_ISL_429639, EPI_ISL_429644, EPI_ISL_429647, EPI_ISL_429648, EPI_ISL_429649, EPI_ISL_429650, EPI_ISL_429651, EPI_ISL_429652, EPI_ISL_429653, EPI_ISL_429654, EPI_ISL_429655 | | | |
| see above | UW Virology Lab | UW Virology Lab | Pavitra Roychoudhury, Hong Xie, Keith Jerome, Alexander Greninger |
| EPI_ISL_430113, EPI_ISL_430114, EPI_ISL_430115, EPI_ISL_430116, EPI_ISL_430117, EPI_ISL_430118, EPI_ISL_430119, EPI_ISL_430120, EPI_ISL_430121, EPI_ISL_430122, EPI_ISL_430123, EPI_ISL_430124, EPI_ISL_430125, EPI_ISL_430126, EPI_ISL_430127, EPI_ISL_430128, EPI_ISL_430129, EPI_ISL_430130, EPI_ISL_430131, EPI_ISL_430132, EPI_ISL_430133, EPI_ISL_430134, EPI_ISL_430135, EPI_ISL_430136, EPI_ISL_430137, EPI_ISL_430138, EPI_ISL_430139, EPI_ISL_430140, EPI_ISL_430141, EPI_ISL_430142, EPI_ISL_430143, EPI_ISL_430144, EPI_ISL_430145, EPI_ISL_430146, EPI_ISL_430147, EPI_ISL_430148, EPI_ISL_430149, EPI_ISL_430150, EPI_ISL_430151, EPI_ISL_430152, EPI_ISL_430153, EPI_ISL_430154, EPI_ISL_430155, EPI_ISL_430156 | | | |
| see above | Seattle Flu Study | Seattle Flu Study | Chu et al |
| EPI_ISL_430160, EPI_ISL_430161, EPI_ISL_430162, EPI_ISL_430163, EPI_ISL_430164, EPI_ISL_430165, EPI_ISL_430166, EPI_ISL_430167, EPI_ISL_430168, EPI_ISL_430169, EPI_ISL_430170, EPI_ISL_430171, EPI_ISL_430172, EPI_ISL_430173, EPI_ISL_430174, EPI_ISL_430175, EPI_ISL_430176, EPI_ISL_430177, EPI_ISL_430178, EPI_ISL_430179, EPI_ISL_430180, EPI_ISL_430181, EPI_ISL_430182, EPI_ISL_430183, EPI_ISL_430184, EPI_ISL_430185, EPI_ISL_430186, EPI_ISL_430187, EPI_ISL_430188, EPI_ISL_430189, EPI_ISL_430190, EPI_ISL_430191, EPI_ISL_430192, EPI_ISL_430193, EPI_ISL_430194, EPI_ISL_430196, EPI_ISL_430197, EPI_ISL_430198, EPI_ISL_430199, EPI_ISL_430200, EPI_ISL_430201, EPI_ISL_430202, EPI_ISL_430203, EPI_ISL_430204, EPI_ISL_430205, EPI_ISL_430206, EPI_ISL_430207, EPI_ISL_430208, EPI_ISL_430209, EPI_ISL_430210, EPI_ISL_430211, EPI_ISL_430212, EPI_ISL_430213, EPI_ISL_430214, EPI_ISL_430215, EPI_ISL_430216, EPI_ISL_430217, EPI_ISL_430218, EPI_ISL_430219, EPI_ISL_430220, EPI_ISL_430221, EPI_ISL_430222, EPI_ISL_430223, EPI_ISL_430224, EPI_ISL_430225, EPI_ISL_430226, EPI_ISL_430227, EPI_ISL_430228, EPI_ISL_430229, EPI_ISL_430230, EPI_ISL_430231, EPI_ISL_430232, EPI_ISL_430233, EPI_ISL_430234, EPI_ISL_430235, EPI_ISL_430236, EPI_ISL_430237, EPI_ISL_430238, EPI_ISL_430239, EPI_ISL_430240, EPI_ISL_430241, EPI_ISL_430242, EPI_ISL_430243, EPI_ISL_430244, EPI_ISL_430245, EPI_ISL_430246, EPI_ISL_430247, EPI_ISL_430248, EPI_ISL_430249, EPI_ISL_430250, EPI_ISL_430251, EPI_ISL_430252, EPI_ISL_430253, EPI_ISL_430254, EPI_ISL_430255, EPI_ISL_430256, EPI_ISL_430257, EPI_ISL_430258, EPI_ISL_430259, EPI_ISL_430260, EPI_ISL_430261, EPI_ISL_430262, EPI_ISL_430263, EPI_ISL_430264, EPI_ISL_430265, EPI_ISL_430266, EPI_ISL_430267, EPI_ISL_430268, EPI_ISL_430269, EPI_ISL_430270, EPI_ISL_430271, EPI_ISL_430272, EPI_ISL_430273, EPI_ISL_430274, EPI_ISL_430275, EPI_ISL_430276, EPI_ISL_430277, EPI_ISL_430278, EPI_ISL_430279, EPI_ISL_430280, EPI_ISL_430281, EPI_ISL_430282, EPI_ISL_430283, EPI_ISL_430284, EPI_ISL_430285, EPI_ISL_430286, EPI_ISL_430287, EPI_ISL_430288, EPI_ISL_430289, EPI_ISL_430290, EPI_ISL_430291, EPI_ISL_430292, EPI_ISL_430293, EPI_ISL_430294, EPI_ISL_430295, EPI_ISL_430296 | | | |
| see above | Washington State Department of Health | Seattle Flu Study | Chu et al |
| EPI_ISL_430867, EPI_ISL_430868, EPI_ISL_430869, EPI_ISL_430870, EPI_ISL_430871, EPI_ISL_430872, EPI_ISL_430873, EPI_ISL_430874, EPI_ISL_430875, EPI_ISL_430876, EPI_ISL_430877, EPI_ISL_430878, EPI_ISL_430880, EPI_ISL_430883, EPI_ISL_430884, EPI_ISL_430885, EPI_ISL_430887, EPI_ISL_430888, EPI_ISL_430889, EPI_ISL_430891, EPI_ISL_430892, EPI_ISL_430893, EPI_ISL_430894, EPI_ISL_430895, EPI_ISL_430896, EPI_ISL_430898, EPI_ISL_430899, EPI_ISL_430900, EPI_ISL_430901, EPI_ISL_430902, EPI_ISL_430904, EPI_ISL_430905, EPI_ISL_430906, EPI_ISL_430911, EPI_ISL_430912, EPI_ISL_430913, EPI_ISL_430914, EPI_ISL_430915, EPI_ISL_430918, EPI_ISL_430920, EPI_ISL_430928, EPI_ISL_430929, EPI_ISL_430930, EPI_ISL_430931, EPI_ISL_430932, EPI_ISL_430933, EPI_ISL_430934, EPI_ISL_430935, EPI_ISL_430936, EPI_ISL_430937, EPI_ISL_430938, EPI_ISL_430939, EPI_ISL_430941, EPI_ISL_430943, EPI_ISL_430951, EPI_ISL_430953, EPI_ISL_430954, EPI_ISL_430955, EPI_ISL_430956, EPI_ISL_430957, EPI_ISL_430958, EPI_ISL_430959, EPI_ISL_430964, EPI_ISL_430966, EPI_ISL_430967, EPI_ISL_430968, EPI_ISL_430969, EPI_ISL_430974, EPI_ISL_430975, EPI_ISL_430976, EPI_ISL_430977, EPI_ISL_430978, EPI_ISL_430979, EPI_ISL_430980 | | | |
| see above | UW Virology Lab | UW Virology Lab | Pavitra Roychoudhury, Hong Xie, Keith Jerome, Alexander Greninger |
| EPI_ISL_434063, EPI_ISL_434064, EPI_ISL_434065, EPI_ISL_434066, EPI_ISL_434067, EPI_ISL_434068, EPI_ISL_434069, EPI_ISL_434070, EPI_ISL_434071, EPI_ISL_434072, EPI_ISL_434073, EPI_ISL_434074, EPI_ISL_434075, EPI_ISL_434076, EPI_ISL_434077, EPI_ISL_434078, EPI_ISL_434079, EPI_ISL_434080, EPI_ISL_434081, EPI_ISL_434082, EPI_ISL_434083, EPI_ISL_434084, EPI_ISL_434085, EPI_ISL_434086, EPI_ISL_434087, EPI_ISL_434088, EPI_ISL_434089, EPI_ISL_434090, EPI_ISL_434091, EPI_ISL_434092, EPI_ISL_434093, EPI_ISL_434094, EPI_ISL_434095, EPI_ISL_434096, EPI_ISL_434097, EPI_ISL_434098, EPI_ISL_434099, EPI_ISL_434100, EPI_ISL_434101, EPI_ISL_434102, EPI_ISL_434103, EPI_ISL_434104, EPI_ISL_434105, EPI_ISL_434106, EPI_ISL_434107, EPI_ISL_434108, EPI_ISL_434109, EPI_ISL_434110, EPI_ISL_434111, EPI_ISL_434112, EPI_ISL_434113, EPI_ISL_434114, EPI_ISL_434115, EPI_ISL_434116, EPI_ISL_434117, EPI_ISL_434118, EPI_ISL_434119, EPI_ISL_434120, EPI_ISL_434121, EPI_ISL_434122, EPI_ISL_434123, EPI_ISL_434124, EPI_ISL_434125, EPI_ISL_434126, EPI_ISL_434127, EPI_ISL_434128, EPI_ISL_434129, EPI_ISL_434130, EPI_ISL_434131, EPI_ISL_434132, EPI_ISL_434133, EPI_ISL_434134, EPI_ISL_434135, EPI_ISL_434136, EPI_ISL_434137, EPI_ISL_434138, EPI_ISL_434139, EPI_ISL_434140, EPI_ISL_434141, EPI_ISL_434142, EPI_ISL_434143, EPI_ISL_434144, EPI_ISL_434145, EPI_ISL_434146, EPI_ISL_434147, EPI_ISL_434148, EPI_ISL_434149, EPI_ISL_434150, EPI_ISL_434151, EPI_ISL_434152, EPI_ISL_434153, EPI_ISL_434154, EPI_ISL_434155, EPI_ISL_434156, EPI_ISL_434157, EPI_ISL_434158, EPI_ISL_434159, EPI_ISL_434160, EPI_ISL_434161, EPI_ISL_434162, EPI_ISL_434163, EPI_ISL_434164, EPI_ISL_434165, EPI_ISL_434166, EPI_ISL_434167, EPI_ISL_434168, EPI_ISL_434169, EPI_ISL_434170, EPI_ISL_434171, EPI_ISL_434172, EPI_ISL_434173, EPI_ISL_434174, EPI_ISL_434175, EPI_ISL_434176, EPI_ISL_434177, EPI_ISL_434178, EPI_ISL_434179, EPI_ISL_434180, EPI_ISL_434181, EPI_ISL_434182, EPI_ISL_434183, EPI_ISL_434184, EPI_ISL_434185, EPI_ISL_434186, EPI_ISL_434187, EPI_ISL_434188, EPI_ISL_434189, EPI_ISL_434190, EPI_ISL_434191, EPI_ISL_434192, EPI_ISL_434193, EPI_ISL_434194, EPI_ISL_434195, EPI_ISL_434196, EPI_ISL_434197, EPI_ISL_434198, EPI_ISL_434199, EPI_ISL_434200, EPI_ISL_434201, EPI_ISL_434202, EPI_ISL_434203, EPI_ISL_434204, EPI_ISL_434205, EPI_ISL_434206, EPI_ISL_434207, EPI_ISL_434208, EPI_ISL_434209, EPI_ISL_434210, EPI_ISL_434211, EPI_ISL_434212, EPI_ISL_434213, EPI_ISL_434214, EPI_ISL_434215, EPI_ISL_434216, EPI_ISL_434217, EPI_ISL_434218, EPI_ISL_434219, EPI_ISL_434220, EPI_ISL_434221, EPI_ISL_434222, EPI_ISL_434223, EPI_ISL_434224, EPI_ISL_434225, EPI_ISL_434226, EPI_ISL_434227, EPI_ISL_434228, EPI_ISL_434229, EPI_ISL_434230, EPI_ISL_434231, EPI_ISL_434232, EPI_ISL_434233, EPI_ISL_434234, EPI_ISL_434235, EPI_ISL_434236, EPI_ISL_434237, EPI_ISL_434238, EPI_ISL_434239, EPI_ISL_434240, EPI_ISL_434241, EPI_ISL_434242, EPI_ISL_434243, EPI_ISL_434244, EPI_ISL_434245, EPI_ISL_434246, EPI_ISL_434247, EPI_ISL_434248, EPI_ISL_434249, EPI_ISL_434250, EPI_ISL_434251, EPI_ISL_434252, EPI_ISL_434253, EPI_ISL_434254, EPI_ISL_434255, EPI_ISL_434256, EPI_ISL_434257, EPI_ISL_434258, EPI_ISL_434259, EPI_ISL_434260, EPI_ISL_434261, EPI_ISL_434262, EPI_ISL_434263, EPI_ISL_434264, EPI_ISL_434265, EPI_ISL_434266, EPI_ISL_434267, EPI_ISL_434268, EPI_ISL_434269, EPI_ISL_434270, EPI_ISL_434271, EPI_ISL_434272, EPI_ISL_434273, EPI_ISL_434274, EPI_ISL_434275, EPI_ISL_434276, EPI_ISL_434277, EPI_ISL_434278, EPI_ISL_434279, EPI_ISL_434280, EPI_ISL_434281, EPI_ISL_434282, EPI_ISL_434283, EPI_ISL_434284, EPI_ISL_434285, EPI_ISL_434286, EPI_ISL_434287, EPI_ISL_434288, EPI_ISL_434289, EPI_ISL_434290, EPI_ISL_434291, EPI_ISL_434292, EPI_ISL_434293, EPI_ISL_434294, EPI_ISL_434295, EPI_ISL_434296, EPI_ISL_434297, EPI_ISL_434298, EPI_ISL_434299, EPI_ISL_434300, EPI_ISL_434301, EPI_ISL_434302, EPI_ISL_434303, EPI_ISL_434304, EPI_ISL_434305, EPI_ISL_434306, EPI_ISL_434307, EPI_ISL_434308, EPI_ISL_434309, EPI_ISL_434310, EPI_ISL_434311, EPI_ISL_434312, EPI_ISL_434313, EPI_ISL_434314, EPI_ISL_434315, EPI_ISL_434316, EPI_ISL_434317, EPI_ISL_434318, EPI_ISL_434319, EPI_ISL_434320, EPI_ISL_434321, EPI_ISL_434322, EPI_ISL_434323, EPI_ISL_434324, EPI_ISL_434325, EPI_ISL_434326, EPI_ISL_434327, EPI_ISL_434328, EPI_ISL_434329, EPI_ISL_434330, EPI_ISL_434331, EPI_ISL_434332, EPI_ISL_434333, EPI_ISL_434334, EPI_ISL_434335, EPI_ISL_434336, EPI_ISL_434337, EPI_ISL_434338, EPI_ISL_434339, EPI_ISL_434340, EPI_ISL_434341, EPI_ISL_434342, EPI_ISL_434343, EPI_ISL_434344, EPI_ISL_434345, EPI_ISL_434346 | | | |
| see above | Washington State Department of Health | Seattle Flu Study | Chu et al |
| EPI_ISL_436040, EPI_ISL_436041, EPI_ISL_436042, EPI_ISL_436043 | DC Public Health Lab Dept of Forensic Science | Pathogen Discovery, Respiratory Viruses Branch, Division of Viral Diseases, Centers for Disease Control and Prevention | Ying Tao, Jing Zhang, Krista Queen, Yan Li, Anna Uehara, Clinton R. Paden, Haibin Wang, Zachary Weiner, Bettina Bankamp, Suxiang Tong |
| EPI_ISL_436612, EPI_ISL_436620 | University of Wisconsin-Madison AIDS Vaccine Research Laboratories | University of Wisconsin-Madison AIDS Vaccine Research Laboratories | Gage Moreno, Katarina Braun, et al. AIDS Vaccine Research Laboratories |
| EPI_ISL_437521, EPI_ISL_437522, EPI_ISL_437527, EPI_ISL_437529, EPI_ISL_437530, EPI_ISL_437531, EPI_ISL_437532, EPI_ISL_437533, EPI_ISL_437534, EPI_ISL_437535 | OHSU Lab Services Molecular Microbiology Lab | Oregon SARS-CoV-2 Genome Sequencing Center | Brendan L. O'Connell, Ruth V. Nichols, Alec J. Hirsch, Guang Fan, Daniel N. Streblow, William B. Messer, Andrew C. Adey, Benjamin N. Bimber, Brian J. O'Roak |
| EPI_ISL_437804, EPI_ISL_437805, EPI_ISL_437806, EPI_ISL_437807, EPI_ISL_437808, EPI_ISL_437809, EPI_ISL_437810, EPI_ISL_437811, EPI_ISL_437812, EPI_ISL_437813, EPI_ISL_437814, EPI_ISL_437815, EPI_ISL_437816, EPI_ISL_437817, EPI_ISL_437820, EPI_ISL_437821, EPI_ISL_437822, EPI_ISL_437823, EPI_ISL_437824, EPI_ISL_437827, EPI_ISL_437829, EPI_ISL_437839, EPI_ISL_437840, EPI_ISL_437841, EPI_ISL_437842, EPI_ISL_437843, EPI_ISL_437844, EPI_ISL_437845, EPI_ISL_437846, EPI_ISL_437847, EPI_ISL_437848, EPI_ISL_437849, EPI_ISL_437850, EPI_ISL_437851, EPI_ISL_437853, EPI_ISL_437854, EPI_ISL_437855, EPI_ISL_437856, EPI_ISL_437857, EPI_ISL_437858, EPI_ISL_437859, EPI_ISL_437860, EPI_ISL_437861, EPI_ISL_437862, EPI_ISL_437863, EPI_ISL_437864 | | | |
| see above | UW Virology Lab | UW Virology Lab | Pavitra Roychoudhury, Hong Xie, Keith Jerome, Alexander Greninger |
| EPI_ISL_438140, EPI_ISL_438141, EPI_ISL_438142, EPI_ISL_438143, EPI_ISL_438144, EPI_ISL_438145, EPI_ISL_438146, EPI_ISL_438147, EPI_ISL_438148, EPI_ISL_438149, EPI_ISL_438150, EPI_ISL_438151, EPI_ISL_438152, EPI_ISL_438153, EPI_ISL_438154, EPI_ISL_438155, EPI_ISL_438156, EPI_ISL_438157, EPI_ISL_438158, EPI_ISL_438159, EPI_ISL_438160, EPI_ISL_438161, EPI_ISL_438162, EPI_ISL_438163, EPI_ISL_438164, EPI_ISL_438165, EPI_ISL_438166, EPI_ISL_438167, EPI_ISL_438168, EPI_ISL_438169, EPI_ISL_438170, EPI_ISL_438171, EPI_ISL_438172, EPI_ISL_438173, EPI_ISL_438174, EPI_ISL_438175 | | | |
| see above | Seattle Flu Study | Seattle Flu Study | Chu et al |
| EPI_ISL_438176, EPI_ISL_438177, EPI_ISL_438178, EPI_ISL_438179, EPI_ISL_438180, EPI_ISL_438181, EPI_ISL_438182, EPI_ISL_438183, EPI_ISL_438184, EPI_ISL_438185, EPI_ISL_438186, EPI_ISL_438187, EPI_ISL_438188, EPI_ISL_438189, EPI_ISL_438190, EPI_ISL_438191, EPI_ISL_438192, EPI_ISL_438193, EPI_ISL_438194, EPI_ISL_438195, EPI_ISL_438196, EPI_ISL_438197, EPI_ISL_438198, EPI_ISL_438199, EPI_ISL_438200, EPI_ISL_438201, EPI_ISL_438202, EPI_ISL_438203, EPI_ISL_438204, EPI_ISL_438205, EPI_ISL_438206, EPI_ISL_438207, EPI_ISL_438208, EPI_ISL_438209, EPI_ISL_438210, EPI_ISL_438211, EPI_ISL_438212, EPI_ISL_438213, EPI_ISL_438214, EPI_ISL_438215, EPI_ISL_438216, EPI_ISL_438217, EPI_ISL_438218, EPI_ISL_438219, EPI_ISL_438220, EPI_ISL_438221 | | | |
| see above | Washington State Department of Health | Seattle Flu Study | Chu et al |
| EPI_ISL_443184, EPI_ISL_443186 | UW Virology Lab | UW Virology Lab | Pavitra Roychoudhury, Hong Xie, Keith Jerome, Alexander Greninger |
| EPI_ISL_444518 | unknown | Molecular Infectious Disease | Anderson,B.P., Rosenthal,S.H., Gerasimova,A., Kagan,R.M. and Owen,R. |
| EPI_ISL_447840 | DC Public Health Lab/ Dept. of Forensic Sciences | Pathogen Discovery, Respiratory Viruses Branch, Division of Viral Diseases, Centers for Disease Control and Prevention | Krista Queen, Yan Li, Anna Uehara, Jing Zhang, Ying Tao, Clinton R. Paden, Haibin Wang, Jasmine Padilla, Mary S. Keckler, Alison S. Laufer Halpin, Justin Lee, Christopher A. Elkins, Suxiang Tong |
| EPI_ISL_449839, EPI_ISL_449840, EPI_ISL_449841, EPI_ISL_449842, EPI_ISL_449843, EPI_ISL_449844, EPI_ISL_449845, EPI_ISL_449846, EPI_ISL_449847, EPI_ISL_449848, EPI_ISL_449849, EPI_ISL_449850, EPI_ISL_449851, EPI_ISL_449852, EPI_ISL_449853, EPI_ISL_449854, EPI_ISL_449855, EPI_ISL_449856, EPI_ISL_449857, EPI_ISL_449858, EPI_ISL_449859, EPI_ISL_449860, EPI_ISL_449861, EPI_ISL_449862, EPI_ISL_449863, EPI_ISL_449864, EPI_ISL_449865, EPI_ISL_449866, EPI_ISL_449867, EPI_ISL_449868, EPI_ISL_449869, EPI_ISL_449870, EPI_ISL_449871, EPI_ISL_449872, EPI_ISL_449873, EPI_ISL_449874, EPI_ISL_449875, EPI_ISL_449876, EPI_ISL_449877, EPI_ISL_449878, EPI_ISL_449879, EPI_ISL_449880, EPI_ISL_449881, EPI_ISL_449882, EPI_ISL_449883, EPI_ISL_449884, EPI_ISL_449885, EPI_ISL_449886, EPI_ISL_449887, EPI_ISL_449888, EPI_ISL_449889, EPI_ISL_449890, EPI_ISL_449891, EPI_ISL_449892, EPI_ISL_449893, EPI_ISL_449894, EPI_ISL_449895, EPI_ISL_449896, EPI_ISL_449897, EPI_ISL_449898, EPI_ISL_449899, EPI_ISL_449900, EPI_ISL_449901, EPI_ISL_449902, EPI_ISL_449903, EPI_ISL_449904, EPI_ISL_449905, EPI_ISL_449906, EPI_ISL_449907, EPI_ISL_449908, EPI_ISL_449909, EPI_ISL_449910, EPI_ISL_449911, EPI_ISL_449912, EPI_ISL_449913, EPI_ISL_449914, EPI_ISL_449915, EPI_ISL_449916, EPI_ISL_449917, EPI_ISL_449918, EPI_ISL_449919, EPI_ISL_449920, EPI_ISL_449921, EPI_ISL_449922, EPI_ISL_449923, EPI_ISL_449924, EPI_ISL_449925, EPI_ISL_449926, EPI_ISL_449927, EPI_ISL_449928, EPI_ISL_449929, EPI_ISL_449930, EPI_ISL_449931, EPI_ISL_449932, EPI_ISL_449933, EPI_ISL_449934, EPI_ISL_449935, EPI_ISL_449936, EPI_ISL_449937, EPI_ISL_449938, EPI_ISL_449939, EPI_ISL_449940, EPI_ISL_449941, EPI_ISL_449942, EPI_ISL_449943, EPI_ISL_449944, EPI_ISL_449945, EPI_ISL_449946, EPI_ISL_449947, EPI_ISL_449948, EPI_ISL_449949, EPI_ISL_449950, EPI_ISL_449951, EPI_ISL_449952, EPI_ISL_449953, EPI_ISL_449954, EPI_ISL_449955, EPI_ISL_449956, EPI_ISL_449957, EPI_ISL_449958, EPI_ISL_449959, EPI_ISL_449960, EPI_ISL_449961, EPI_ISL_449962, EPI_ISL_449963, EPI_ISL_449964, EPI_ISL_449965, EPI_ISL_449966, EPI_ISL_449967, EPI_ISL_449968, EPI_ISL_449969, EPI_ISL_449970, EPI_ISL_449971, EPI_ISL_449972, EPI_ISL_449973, EPI_ISL_449974, EPI_ISL_449975, EPI_ISL_449976, EPI_ISL_449977, EPI_ISL_449978, EPI_ISL_449979, EPI_ISL_449980, EPI_ISL_449981, EPI_ISL_449982, EPI_ISL_449983, EPI_ISL_449984, EPI_ISL_449985, EPI_ISL_449986, EPI_ISL_449987, EPI_ISL_449988, EPI_ISL_449989, EPI_ISL_449990, EPI_ISL_449991, EPI_ISL_449992, EPI_ISL_449993, EPI_ISL_449994, EPI_ISL_449995, EPI_ISL_449996, EPI_ISL_449997, EPI_ISL_449998, EPI_ISL_449999, EPI_ISL_450000, EPI_ISL_450001, EPI_ISL_450002, EPI_ISL_450003 | | | |
| see above | Washington State Department of | Seattle Flu Study | Chu et al |

| | | | |
|---|---|---|---|
| | Health | | |
| EPI_ISL_454692 | Quest Diagnostics | Quest Diagnostics | Anderson,B.P., Rosenthal,S.H., Gerasimova,A., Kagan,R.M. and Owen, R. |
| EPI_ISL_457753, EPI_ISL_457757, EPI_ISL_457760, EPI_ISL_457770, EPI_ISL_457771, EPI_ISL_457775, EPI_ISL_457776, EPI_ISL_457781, EPI_ISL_457793, EPI_ISL_457796, EPI_ISL_457797, EPI_ISL_457801, EPI_ISL_457805, EPI_ISL_457809, EPI_ISL_457810, EPI_ISL_457814, EPI_ISL_457822, EPI_ISL_457823 | | | |
| see above | Johns Hopkins Hospital Department of Pathology | Johns Hopkins Hospital Department of Pathology | Peter M. Thielen, Thomas Mehoke, Shirlee Wohl, Srividya Ramakrishnan, Melanie Kirsche, Amanda Ernlund, Craig Howser, Kristina Zudock, Oluwaseun Falade-Nwulia, Norah Sadowski, Paul Morris, Mark Hopkins, Yunfan Fan, Nidia Trovao, Victoria Gniazdowski, Michael C. Schatz, Stuart C. Ray, Winston Timp, Heba H. Mostafa |
| EPI_ISL_460621, EPI_ISL_460623, EPI_ISL_460624, EPI_ISL_460625, EPI_ISL_460626, EPI_ISL_460627, EPI_ISL_460630, EPI_ISL_460631, EPI_ISL_460632, EPI_ISL_460633, EPI_ISL_460634, EPI_ISL_461399, EPI_ISL_461400, EPI_ISL_461401, EPI_ISL_461402, EPI_ISL_461403, EPI_ISL_461404, EPI_ISL_461405, EPI_ISL_461406, EPI_ISL_461407, EPI_ISL_461410, EPI_ISL_461413, EPI_ISL_461415, EPI_ISL_461418, EPI_ISL_461419, EPI_ISL_461421, EPI_ISL_461422, EPI_ISL_461423, EPI_ISL_461426, EPI_ISL_461427, EPI_ISL_461428, EPI_ISL_461429, EPI_ISL_461430, EPI_ISL_461431, EPI_ISL_461432, EPI_ISL_461433, EPI_ISL_461434, EPI_ISL_461436, EPI_ISL_461439, EPI_ISL_461440, EPI_ISL_461442, EPI_ISL_461443, EPI_ISL_461444, EPI_ISL_461445, EPI_ISL_461446, EPI_ISL_461447, EPI_ISL_461448, EPI_ISL_461449, EPI_ISL_461450, EPI_ISL_461451, EPI_ISL_461452, EPI_ISL_461453, EPI_ISL_461454, EPI_ISL_461455, EPI_ISL_461456, EPI_ISL_461457, EPI_ISL_461458, EPI_ISL_461459, EPI_ISL_461460, EPI_ISL_461461, EPI_ISL_461462, EPI_ISL_461463, EPI_ISL_461464, EPI_ISL_461465, EPI_ISL_461466, EPI_ISL_461467, EPI_ISL_461468, EPI_ISL_461469, EPI_ISL_461470, EPI_ISL_461471, EPI_ISL_461472, EPI_ISL_461473, EPI_ISL_461474, EPI_ISL_461475, EPI_ISL_461476, EPI_ISL_461477 | | | |
| see above | UW Virology Lab | UW Virology Lab | Pavitra Roychoudhury, Amin Addetia, Hong Xie, Lasata Shrestha, Truong Nguyen, Meei-Li Huang, Keith Jerome, Alexander Greninger |
| EPI_ISL_463324, EPI_ISL_463325, EPI_ISL_463326, EPI_ISL_463327, EPI_ISL_463328, EPI_ISL_463329, EPI_ISL_463330, EPI_ISL_463331, EPI_ISL_463332, EPI_ISL_463333, EPI_ISL_463334, EPI_ISL_463335, EPI_ISL_463336, EPI_ISL_463337, EPI_ISL_463338, EPI_ISL_463339, EPI_ISL_463340, EPI_ISL_463341, EPI_ISL_463342, EPI_ISL_463343, EPI_ISL_463344, EPI_ISL_463345, EPI_ISL_463346, EPI_ISL_463347, EPI_ISL_463348, EPI_ISL_463349, EPI_ISL_463350, EPI_ISL_463351, EPI_ISL_463352, EPI_ISL_463353, EPI_ISL_463354, EPI_ISL_463355, EPI_ISL_463356, EPI_ISL_463357, EPI_ISL_463358, EPI_ISL_463359, EPI_ISL_463360, EPI_ISL_463361, EPI_ISL_463362, EPI_ISL_463363, EPI_ISL_463364, EPI_ISL_463365, EPI_ISL_463366, EPI_ISL_463367, EPI_ISL_463368, EPI_ISL_463369, EPI_ISL_463370, EPI_ISL_463371, EPI_ISL_463372, EPI_ISL_463373, EPI_ISL_463374, EPI_ISL_463375, EPI_ISL_463376, EPI_ISL_463377, EPI_ISL_463378, EPI_ISL_463379, EPI_ISL_463380, EPI_ISL_463381, EPI_ISL_463382, EPI_ISL_463383, EPI_ISL_463384, EPI_ISL_463385, EPI_ISL_463386, EPI_ISL_463387, EPI_ISL_463388, EPI_ISL_463389, EPI_ISL_463390, EPI_ISL_463391, EPI_ISL_463392, EPI_ISL_463393, EPI_ISL_463394, EPI_ISL_463395, EPI_ISL_463396, EPI_ISL_463397, EPI_ISL_463398, EPI_ISL_463399, EPI_ISL_463400, EPI_ISL_463401, EPI_ISL_463402, EPI_ISL_463403, EPI_ISL_463404, EPI_ISL_463405, EPI_ISL_463406, EPI_ISL_463407, EPI_ISL_463408, EPI_ISL_463409, EPI_ISL_463410, EPI_ISL_463411, EPI_ISL_463412, EPI_ISL_463413, EPI_ISL_463414, EPI_ISL_463415, EPI_ISL_463416, EPI_ISL_463417, EPI_ISL_463418, EPI_ISL_463419, EPI_ISL_463420, EPI_ISL_463421, EPI_ISL_463422, EPI_ISL_463423, EPI_ISL_463424, EPI_ISL_463425, EPI_ISL_463426, EPI_ISL_463427, EPI_ISL_463428, EPI_ISL_463429, EPI_ISL_463430, EPI_ISL_463431, EPI_ISL_463432, EPI_ISL_463433, EPI_ISL_463434, EPI_ISL_463435, EPI_ISL_463436, EPI_ISL_463437, EPI_ISL_463438, EPI_ISL_463439, EPI_ISL_463440, EPI_ISL_463441, EPI_ISL_463442, EPI_ISL_463443, EPI_ISL_463444, EPI_ISL_463445, EPI_ISL_463446, EPI_ISL_463447, EPI_ISL_463448, EPI_ISL_463449, EPI_ISL_463450, EPI_ISL_463451, EPI_ISL_463452, EPI_ISL_463453, EPI_ISL_463454, EPI_ISL_463455, EPI_ISL_463456, EPI_ISL_463457, EPI_ISL_463458, EPI_ISL_463459, EPI_ISL_463460, EPI_ISL_463461, EPI_ISL_463462, EPI_ISL_463463, EPI_ISL_463464, EPI_ISL_463465, EPI_ISL_463466, EPI_ISL_463467, EPI_ISL_463468, EPI_ISL_463469, EPI_ISL_463470, EPI_ISL_463471, EPI_ISL_463472, EPI_ISL_463473, EPI_ISL_463474, EPI_ISL_463475, EPI_ISL_463476, EPI_ISL_463477, EPI_ISL_463478, EPI_ISL_463479, EPI_ISL_463480, EPI_ISL_463481, EPI_ISL_463482, EPI_ISL_463483, EPI_ISL_463484, EPI_ISL_463485, EPI_ISL_463486, EPI_ISL_463487, EPI_ISL_463488, EPI_ISL_463489, EPI_ISL_463490, EPI_ISL_463491, EPI_ISL_463492, EPI_ISL_463493, EPI_ISL_463494, EPI_ISL_463495, EPI_ISL_463496, EPI_ISL_463497, EPI_ISL_463498, EPI_ISL_463499, EPI_ISL_463500, EPI_ISL_463501, EPI_ISL_463502, EPI_ISL_463503, EPI_ISL_463504, EPI_ISL_463505, EPI_ISL_463506, EPI_ISL_463507, EPI_ISL_463508, EPI_ISL_463509, EPI_ISL_463510, EPI_ISL_463511, EPI_ISL_463512, EPI_ISL_463513, EPI_ISL_463514, EPI_ISL_463515, EPI_ISL_463516, EPI_ISL_463517, EPI_ISL_463518, EPI_ISL_463519, EPI_ISL_463520, EPI_ISL_463521, EPI_ISL_463522, EPI_ISL_463523, EPI_ISL_463524, EPI_ISL_463525, EPI_ISL_463526, EPI_ISL_463527, EPI_ISL_463528, EPI_ISL_463529, EPI_ISL_463530, EPI_ISL_463531, EPI_ISL_463532, EPI_ISL_463533, EPI_ISL_463534, EPI_ISL_463535, EPI_ISL_463536, EPI_ISL_463537, EPI_ISL_463538, EPI_ISL_463539, EPI_ISL_463540, EPI_ISL_463541, EPI_ISL_463542, EPI_ISL_463543, EPI_ISL_463544, EPI_ISL_463545, EPI_ISL_463546, EPI_ISL_463547, EPI_ISL_463548, EPI_ISL_463549, EPI_ISL_463550, EPI_ISL_463551, EPI_ISL_463552, EPI_ISL_463553, EPI_ISL_463554, EPI_ISL_463555, EPI_ISL_463556, EPI_ISL_463557, EPI_ISL_463558, EPI_ISL_463559, EPI_ISL_463560, EPI_ISL_463561, EPI_ISL_463562, EPI_ISL_463563, EPI_ISL_463564, EPI_ISL_463565, EPI_ISL_463566, EPI_ISL_463567, EPI_ISL_463568, EPI_ISL_463569, EPI_ISL_463570, EPI_ISL_463571, EPI_ISL_463572, EPI_ISL_463573, EPI_ISL_463574, EPI_ISL_463575, EPI_ISL_463576, EPI_ISL_463577, EPI_ISL_463578, EPI_ISL_463579, EPI_ISL_463580, EPI_ISL_463581, EPI_ISL_463582, EPI_ISL_463583, EPI_ISL_463584, EPI_ISL_463585, EPI_ISL_463586, EPI_ISL_463587, EPI_ISL_463588, EPI_ISL_463589, EPI_ISL_463590, EPI_ISL_463591, EPI_ISL_463592, EPI_ISL_463593, EPI_ISL_463594, EPI_ISL_463595, EPI_ISL_463596, EPI_ISL_463597, EPI_ISL_463598, EPI_ISL_463599, EPI_ISL_463600, EPI_ISL_463601, EPI_ISL_463602, EPI_ISL_463603, EPI_ISL_463604, EPI_ISL_463605, EPI_ISL_463606, EPI_ISL_463607, EPI_ISL_463608, EPI_ISL_463609, EPI_ISL_463610, EPI_ISL_463611, EPI_ISL_463612, EPI_ISL_463613, EPI_ISL_463614, EPI_ISL_463615, EPI_ISL_463616, EPI_ISL_463617, EPI_ISL_463618, EPI_ISL_463619, EPI_ISL_463620, EPI_ISL_463621, EPI_ISL_463622, EPI_ISL_463623, EPI_ISL_463624, EPI_ISL_463625, EPI_ISL_463626, EPI_ISL_463627, EPI_ISL_463628, EPI_ISL_463629, EPI_ISL_463630, EPI_ISL_463631, EPI_ISL_463632, EPI_ISL_463633, EPI_ISL_463634, EPI_ISL_463635, EPI_ISL_463636, EPI_ISL_463637, EPI_ISL_463638, EPI_ISL_463639, EPI_ISL_463640, EPI_ISL_463641, EPI_ISL_463642, EPI_ISL_463643, EPI_ISL_463644, EPI_ISL_463645, EPI_ISL_463646, EPI_ISL_463647, EPI_ISL_463648, EPI_ISL_463649, EPI_ISL_463650, EPI_ISL_463651, EPI_ISL_463652, EPI_ISL_463653, EPI_ISL_463654, EPI_ISL_463655, EPI_ISL_463656, EPI_ISL_463657, EPI_ISL_463658, EPI_ISL_463659, EPI_ISL_463660, EPI_ISL_463661, EPI_ISL_463662, EPI_ISL_463663, EPI_ISL_463664, EPI_ISL_463665, EPI_ISL_463666, EPI_ISL_463667, EPI_ISL_463668, EPI_ISL_463669, EPI_ISL_463670, EPI_ISL_463671, EPI_ISL_463672, EPI_ISL_463673, EPI_ISL_463674, EPI_ISL_463675, EPI_ISL_463676, EPI_ISL_463677, EPI_ISL_463678, EPI_ISL_463679, EPI_ISL_463680, EPI_ISL_463681, EPI_ISL_463682, EPI_ISL_463683, EPI_ISL_463684, EPI_ISL_463685, EPI_ISL_463686, EPI_ISL_463687, EPI_ISL_463688, EPI_ISL_463689, EPI_ISL_463690, EPI_ISL_463691, EPI_ISL_463692, EPI_ISL_463693, EPI_ISL_463694, EPI_ISL_463695, EPI_ISL_463696, EPI_ISL_463697, EPI_ISL_463698, EPI_ISL_463699, EPI_ISL_463700, EPI_ISL_463701, EPI_ISL_463702, EPI_ISL_463703, EPI_ISL_463704, EPI_ISL_463705 | | | |
| see above | Washington State Department of Health | Seattle Flu Study | Chu et al |
| EPI_ISL_467926, EPI_ISL_467927 | Quest Diagnostics | Quest Diagnostics | Anderson,B.P., Rosenthal,S.H., Gerasimova,A., Kagan,R.M. and Owen, R. |
| EPI_ISL_468080, EPI_ISL_468081, EPI_ISL_468082, EPI_ISL_468083, EPI_ISL_468084, EPI_ISL_468086, EPI_ISL_468087, EPI_ISL_468088, EPI_ISL_468089, EPI_ISL_468093, EPI_ISL_468094, EPI_ISL_468095, EPI_ISL_468098, EPI_ISL_468099, EPI_ISL_468103, EPI_ISL_468104, EPI_ISL_468106, EPI_ISL_468110, EPI_ISL_468111, EPI_ISL_468112, EPI_ISL_468113, EPI_ISL_468115, EPI_ISL_468116, EPI_ISL_468117, EPI_ISL_468118 | | | |
| see above | OHSU Lab Services Molecular Microbiology Lab | Oregon SARS-CoV-2 Genome Sequencing Center | Brendan L. O'Connell, Ruth V. Nichols, Alec J. Hirsch, Guang Fan, Daniel N. Streblow, William B. Messer, Andrew C. Adey, Benjamin N. Bimber, Brian J. O'Roak |
| EPI_ISL_468590 | Quest Diagnostics | Quest Diagnostics | Anderson,B.P., Rosenthal,S.H., Gerasimova,A., Kagan,R.M. and Owen, R. |
| EPI_ISL_476900, EPI_ISL_476901, EPI_ISL_476902, EPI_ISL_476903, EPI_ISL_476905, EPI_ISL_476906, EPI_ISL_476907, EPI_ISL_476908, EPI_ISL_476911, EPI_ISL_476920, EPI_ISL_476921, EPI_ISL_476924, EPI_ISL_476925, EPI_ISL_476927, EPI_ISL_476928, EPI_ISL_476929, EPI_ISL_476930, EPI_ISL_476933, EPI_ISL_476934, EPI_ISL_476935, EPI_ISL_476936, EPI_ISL_476937, EPI_ISL_476938, EPI_ISL_476939, EPI_ISL_476940, EPI_ISL_477685, EPI_ISL_477686, EPI_ISL_477687, EPI_ISL_477688, EPI_ISL_477689, EPI_ISL_477690, EPI_ISL_477691, EPI_ISL_477692, EPI_ISL_477693, EPI_ISL_477694, EPI_ISL_477695, EPI_ISL_477696, EPI_ISL_477697, EPI_ISL_477698, EPI_ISL_477699, EPI_ISL_477700, EPI_ISL_477701, EPI_ISL_477702, EPI_ISL_477703, EPI_ISL_477704, EPI_ISL_477705, EPI_ISL_477706, EPI_ISL_477707, EPI_ISL_477708, EPI_ISL_477709, EPI_ISL_477710, EPI_ISL_477711, EPI_ISL_477712, EPI_ISL_477713, EPI_ISL_477714, EPI_ISL_477715, EPI_ISL_477716, EPI_ISL_477717, EPI_ISL_477718, EPI_ISL_477719, EPI_ISL_477720, EPI_ISL_477721, EPI_ISL_477722, EPI_ISL_477723, EPI_ISL_477724 | | | |
| see above | UW Virology Lab | UW Virology Lab | Pavitra Roychoudhury, Hong Xie, Lasata Shrestha, Amin Addetia, Truong Nguyen, Victoria M Rachleff, Meei-Li Huang, Keith R Jerome, Alexander Greninger |
| EPI_ISL_480395, EPI_ISL_480397, EPI_ISL_480398, EPI_ISL_480410 | University of Wisconsin-Madison AIDS Vaccine Research Laboratories | University of Wisconsin-Madison AIDS Vaccine Research Laboratories | Gage Moreno, Katarina Braun, et al. AIDS Vaccine Research Laboratories |
| EPI_ISL_485971, EPI_ISL_485972, EPI_ISL_485973, EPI_ISL_485974, EPI_ISL_485975, EPI_ISL_485976, EPI_ISL_485977, EPI_ISL_485978, EPI_ISL_485979, EPI_ISL_485980, EPI_ISL_485981, EPI_ISL_485982, EPI_ISL_485983, EPI_ISL_485984, EPI_ISL_485985, EPI_ISL_485986, EPI_ISL_485987, EPI_ISL_485988, EPI_ISL_485989, EPI_ISL_485990, EPI_ISL_485991, EPI_ISL_485992, EPI_ISL_485993, EPI_ISL_485994, EPI_ISL_485995, EPI_ISL_485996, EPI_ISL_485997, EPI_ISL_485998, EPI_ISL_485999, EPI_ISL_486000, EPI_ISL_486001, EPI_ISL_486002, EPI_ISL_486003, EPI_ISL_486004, EPI_ISL_486005, EPI_ISL_486006, EPI_ISL_486007, EPI_ISL_486008, EPI_ISL_486009, EPI_ISL_486010, EPI_ISL_486011, EPI_ISL_486012, EPI_ISL_486013, EPI_ISL_486014, EPI_ISL_486015, EPI_ISL_486016, EPI_ISL_486017, EPI_ISL_486018, EPI_ISL_486019, EPI_ISL_486020, EPI_ISL_486021, EPI_ISL_486022, EPI_ISL_486023, EPI_ISL_486024, EPI_ISL_486025, EPI_ISL_486026, EPI_ISL_486027, EPI_ISL_486028, EPI_ISL_486029, EPI_ISL_486030, EPI_ISL_486032, EPI_ISL_486033, EPI_ISL_486034, EPI_ISL_486035, EPI_ISL_486036, EPI_ISL_486037, EPI_ISL_486038, EPI_ISL_486039, EPI_ISL_486040, EPI_ISL_486041, EPI_ISL_486042, EPI_ISL_486043, EPI_ISL_486044, EPI_ISL_486045, EPI_ISL_486046, EPI_ISL_486047, EPI_ISL_486048, EPI_ISL_486049, EPI_ISL_486050, EPI_ISL_486051, EPI_ISL_486052, EPI_ISL_486053, EPI_ISL_486054, EPI_ISL_486055, EPI_ISL_486056, EPI_ISL_486057, EPI_ISL_486058, EPI_ISL_486059, EPI_ISL_486060, EPI_ISL_486061, EPI_ISL_486062, EPI_ISL_486063, EPI_ISL_486064, EPI_ISL_486065, EPI_ISL_486066, EPI_ISL_486067, EPI_ISL_486068, EPI_ISL_486069, EPI_ISL_486070, EPI_ISL_486071, EPI_ISL_486072, EPI_ISL_486073, EPI_ISL_486074, EPI_ISL_486075, EPI_ISL_486076, EPI_ISL_486077, EPI_ISL_486078, EPI_ISL_486079, EPI_ISL_486080, EPI_ISL_486081, EPI_ISL_486082, EPI_ISL_486083, EPI_ISL_486084, EPI_ISL_486085, EPI_ISL_486086, EPI_ISL_486087, EPI_ISL_486088, EPI_ISL_486089, EPI_ISL_486090, EPI_ISL_486093, EPI_ISL_486094, EPI_ISL_486095, EPI_ISL_486097, EPI_ISL_486098, EPI_ISL_486099, EPI_ISL_486102, EPI_ISL_486103, EPI_ISL_486105, EPI_ISL_486106, EPI_ISL_486108, EPI_ISL_486109, EPI_ISL_486110, EPI_ISL_486112, EPI_ISL_486113, EPI_ISL_486114, EPI_ISL_491005, EPI_ISL_491006, EPI_ISL_491007, EPI_ISL_491009, EPI_ISL_491010, EPI_ISL_491011, EPI_ISL_491012, EPI_ISL_491013, EPI_ISL_491014, EPI_ISL_491015, EPI_ISL_491016, EPI_ISL_491017, EPI_ISL_491018, EPI_ISL_491019, EPI_ISL_491020, EPI_ISL_491021, EPI_ISL_491022, EPI_ISL_491023, EPI_ISL_491024, EPI_ISL_491025, EPI_ISL_491026, EPI_ISL_491027, EPI_ISL_491028, EPI_ISL_491029, EPI_ISL_491030, EPI_ISL_491031, EPI_ISL_491032, EPI_ISL_491033, EPI_ISL_491034 | | | |
| see above | UW Virology Lab | UW Virology Lab | Pavitra Roychoudhury, Hong Xie, Lasata Shrestha, Amin Addetia, Truong Nguyen, Victoria M Rachleff, Meei-Li Huang, Keith R Jerome, Alexander Greninger |
| EPI_ISL_495642, EPI_ISL_495643, EPI_ISL_495644, EPI_ISL_495645, EPI_ISL_495646, EPI_ISL_495647, EPI_ISL_495648, EPI_ISL_495649, EPI_ISL_495650, EPI_ISL_495651, EPI_ISL_495652, EPI_ISL_495653 | | | |
| see above | Washington State Department of Health | Seattle Flu Study | Deborah A. Nickerson, Chris D. Frazar, Jover Lee, Benjamin Pelle, Matthew Richardson, Amanda Adler, Elisabeth Brandstetter, Peter D. Han, Kairsten Fay, Misja Ilcisin, Kirsten Lacombe, Thomas R. Sibley, Melissa Truong, Caitlin R. Wolf, Romesh Gautom, Geoff Melly, Brian Hiatt, Philip Dykema, Scott Lindquist, Michael Boeckh, Janet A. Englund, Michael Famulare, Barry R. Lutz, Mark J. Rieder, Lea M. Starita, Matthew Thompson, Helen Y. Chu, Jay Shendure, Trevor Bedford |
| EPI_ISL_495654, EPI_ISL_495655, EPI_ISL_495656, EPI_ISL_495657, EPI_ISL_495658, EPI_ISL_495659, EPI_ISL_495660, EPI_ISL_495661, EPI_ISL_495662 | Seattle Flu Study | Seattle Flu Study | Deborah A. Nickerson, Chris D. Frazar, Jover Lee, Benjamin Pelle, Matthew Richardson, Amanda Adler, Elisabeth Brandstetter, Peter D. Han, Kairsten Fay, Misja Ilcisin, Kirsten Lacombe, Thomas R. Sibley, Melissa Truong, Caitlin R. Wolf, Karen Cowgill, Stephanie Schrag, Jeff Duchin, Michael Boeckh, Janet A. Englund, Michael Famulare, Barry R. Lutz, Mark J. Rieder, Lea M. Starita, Matthew Thompson, Helen Y. Chu, Trevor Bedford, Jay Shendure |
| EPI_ISL_495663 | Seattle Flu Study | Seattle Flu Study | Deborah A. Nickerson, Chris D. Frazar, Jover Lee, Benjamin Pelle, Matthew Richardson, Amanda Adler, Elisabeth Brandstetter, Peter D. Han, Kairsten Fay, Misja Ilcisin, Kirsten Lacombe, Thomas R. Sibley, Melissa Truong, Caitlin R. Wolf, Michael Boeckh, Janet A. Englund, Michael Famulare, Barry R. Lutz, Mark J. Rieder, Lea M. Starita, Matthew Thompson, Jay Shendure, Trevor Bedford, Helen Y. Chu |
| EPI_ISL_495664, EPI_ISL_495665 | Seattle Flu Study | Seattle Flu Study | Deborah A. Nickerson, Chris D. Frazar, Jover Lee, Benjamin Pelle, Matthew Richardson, Amanda Adler, Elisabeth Brandstetter, Peter D. Han, Kairsten Fay, Misja Ilcisin, Kirsten Lacombe, Thomas R. Sibley, Melissa Truong, Caitlin R. Wolf, Karen Cowgill, Stephanie Schrag, Jeff Duchin, Michael Boeckh, Janet A. Englund, Michael Famulare, Barry R. Lutz, Mark J. Rieder, Lea M. Starita, Matthew Thompson, Helen Y. Chu, Trevor Bedford, Jay Shendure |
| EPI_ISL_495666, EPI_ISL_495667, EPI_ISL_495668, EPI_ISL_495669, EPI_ISL_495670, EPI_ISL_495671, EPI_ISL_495672, EPI_ISL_495673, EPI_ISL_495674, EPI_ISL_495675, EPI_ISL_495676, EPI_ISL_495677, EPI_ISL_495678, EPI_ISL_495679, EPI_ISL_495680, EPI_ISL_495681, EPI_ISL_495682, EPI_ISL_495683, EPI_ISL_495684, EPI_ISL_495685, EPI_ISL_495686, EPI_ISL_495687, EPI_ISL_495688, EPI_ISL_495689, EPI_ISL_495690, EPI_ISL_495691, EPI_ISL_495692, EPI_ISL_495693, EPI_ISL_495694, EPI_ISL_495695, EPI_ISL_495696, EPI_ISL_495697, EPI_ISL_495698, EPI_ISL_495699, EPI_ISL_495700, EPI_ISL_495701, EPI_ISL_495702, EPI_ISL_495703, EPI_ISL_495704, EPI_ISL_495705, EPI_ISL_495706, EPI_ISL_495707, EPI_ISL_495708, EPI_ISL_495709, EPI_ISL_495710, EPI_ISL_495711, EPI_ISL_495712, EPI_ISL_495713, EPI_ISL_495714, EPI_ISL_495715, EPI_ISL_495716, EPI_ISL_495717, EPI_ISL_495718, EPI_ISL_495719, EPI_ISL_495720, EPI_ISL_495721, EPI_ISL_495722, EPI_ISL_495723, EPI_ISL_495724, EPI_ISL_495725, EPI_ISL_495726, EPI_ISL_495727, EPI_ISL_495728, EPI_ISL_495729, EPI_ISL_495730, EPI_ISL_495731, EPI_ISL_495732, EPI_ISL_495733, EPI_ISL_495734, EPI_ISL_495735, EPI_ISL_495736, EPI_ISL_495737, EPI_ISL_495738, EPI_ISL_495739, EPI_ISL_495740, EPI_ISL_495741, EPI_ISL_495742, EPI_ISL_495743, EPI_ISL_495744, EPI_ISL_495745, EPI_ISL_495746, EPI_ISL_495747, EPI_ISL_495748, EPI_ISL_495749, EPI_ISL_495750, EPI_ISL_495751, EPI_ISL_495752, EPI_ISL_495753, EPI_ISL_495754, EPI_ISL_495755, EPI_ISL_495756, EPI_ISL_495757, EPI_ISL_495758, EPI_ISL_495759, EPI_ISL_495760, EPI_ISL_495761, EPI_ISL_495762, EPI_ISL_495763, EPI_ISL_495764, EPI_ISL_495765, EPI_ISL_495766, EPI_ISL_495767, EPI_ISL_495768, EPI_ISL_495769, EPI_ISL_495770, EPI_ISL_495771, EPI_ISL_495772, EPI_ISL_495773, EPI_ISL_495774, EPI_ISL_495775, EPI_ISL_495776, EPI_ISL_495777, EPI_ISL_495778, EPI_ISL_495779, EPI_ISL_495780, EPI_ISL_495781, EPI_ISL_495782, EPI_ISL_495783, EPI_ISL_495784, EPI_ISL_495785, EPI_ISL_495786, EPI_ISL_495787, EPI_ISL_495788, EPI_ISL_495789, EPI_ISL_495790, EPI_ISL_495791, EPI_ISL_495792, EPI_ISL_495793, EPI_ISL_495794, EPI_ISL_495795, EPI_ISL_495796, EPI_ISL_495797, EPI_ISL_495798, EPI_ISL_495799, EPI_ISL_495800, EPI_ISL_495801, EPI_ISL_495802, EPI_ISL_495803, EPI_ISL_495804, EPI_ISL_495805, EPI_ISL_495806, EPI_ISL_495807, EPI_ISL_495808, EPI_ISL_495809, EPI_ISL_495810, EPI_ISL_495811, EPI_ISL_495812, EPI_ISL_495813, EPI_ISL_495814, EPI_ISL_495815, EPI_ISL_495816, EPI_ISL_495817, EPI_ISL_495818, EPI_ISL_495819, EPI_ISL_495820, EPI_ISL_495821, EPI_ISL_495822, EPI_ISL_495823, EPI_ISL_495824, EPI_ISL_495825, EPI_ISL_495826, EPI_ISL_495827, EPI_ISL_495828, EPI_ISL_495829, EPI_ISL_495830, EPI_ISL_495831, EPI_ISL_495832, EPI_ISL_495833, EPI_ISL_495834, EPI_ISL_495835, EPI_ISL_495836, EPI_ISL_495837, EPI_ISL_495838, EPI_ISL_495839, EPI_ISL_495840, EPI_ISL_495841, EPI_ISL_495842, EPI_ISL_495843, EPI_ISL_495844, EPI_ISL_495845, EPI_ISL_495846, EPI_ISL_495847, EPI_ISL_495848, EPI_ISL_495849, EPI_ISL_495850, EPI_ISL_495851, EPI_ISL_495852, EPI_ISL_495853, EPI_ISL_495854, EPI_ISL_495855, EPI_ISL_495856, EPI_ISL_495857, EPI_ISL_495858, EPI_ISL_495859, EPI_ISL_495860, EPI_ISL_495861, EPI_ISL_495862, EPI_ISL_495863, EPI_ISL_495864, EPI_ISL_495865, EPI_ISL_495866, EPI_ISL_495867, EPI_ISL_495868, EPI_ISL_495869, EPI_ISL_495870, EPI_ISL_495871, EPI_ISL_495872, EPI_ISL_495873, EPI_ISL_495874, EPI_ISL_495875, EPI_ISL_495876, EPI_ISL_495877, EPI_ISL_495878, EPI_ISL_495879, EPI_ISL_495880, EPI_ISL_495881, EPI_ISL_495882, EPI_ISL_495883, EPI_ISL_495884, EPI_ISL_495885, EPI_ISL_495886, EPI_ISL_495887, EPI_ISL_495888, EPI_ISL_495889, EPI_ISL_495890, EPI_ISL_495891, EPI_ISL_495892, EPI_ISL_495893, EPI_ISL_495894, EPI_ISL_495895, EPI_ISL_495896, EPI_ISL_495897, EPI_ISL_495898, EPI_ISL_495899, EPI_ISL_495900, EPI_ISL_495901, EPI_ISL_495902, EPI_ISL_495903, EPI_ISL_495904, EPI_ISL_495905, EPI_ISL_495906, EPI_ISL_495907, EPI_ISL_495908, EPI_ISL_495909, EPI_ISL_495910, EPI_ISL_495911, EPI_ISL_495912, EPI_ISL_495913, EPI_ISL_495914, EPI_ISL_495915, EPI_ISL_495916 | | | |
| see above | Washington State Department of Health | Seattle Flu Study | Deborah A. Nickerson, Chris D. Frazar, Jover Lee, Benjamin Pelle, Matthew Richardson, Amanda Adler, Elisabeth Brandstetter, Peter D. Han, Kairsten Fay, Misja Ilcisin, Kirsten Lacombe, Thomas R. Sibley, Melissa Truong, Caitlin R. Wolf, Romesh Gautom, Geoff |
| EPI_ISL_495917, EPI_ISL_495918, EPI_ISL_495919, EPI_ISL_495920, EPI_ISL_495921, EPI_ISL_495922, EPI_ISL_495923, EPI_ISL_495924, EPI_ISL_495925, EPI_ISL_495926, EPI_ISL_495927, EPI_ISL_495928, EPI_ISL_495929, EPI_ISL_495930, EPI_ISL_495931, EPI_ISL_495932, EPI_ISL_495933, EPI_ISL_495934, EPI_ISL_495935, EPI_ISL_495936, EPI_ISL_495937, EPI_ISL_495938, EPI_ISL_495939, EPI_ISL_495940, EPI_ISL_495941, EPI_ISL_495942, EPI_ISL_495943, EPI_ISL_495944, EPI_ISL_495945, EPI_ISL_495946, EPI_ISL_495947, EPI_ISL_495948, EPI_ISL_495949, EPI_ISL_495950, EPI_ISL_495951, EPI_ISL_495952, EPI_ISL_495953, EPI_ISL_495954, EPI_ISL_495955, EPI_ISL_495956, EPI_ISL_495957, EPI_ISL_495958, EPI_ISL_495959, EPI_ISL_495960, EPI_ISL_495961, EPI_ISL_495962, EPI_ISL_495963, EPI_ISL_495964, EPI_ISL_495965, EPI_ISL_495966, EPI_ISL_495967, EPI_ISL_495968, EPI_ISL_495969, EPI_ISL_495970, EPI_ISL_495971, EPI_ISL_495972, EPI_ISL_495973, EPI_ISL_495974, EPI_ISL_495975, EPI_ISL_495976, EPI_ISL_495977, EPI_ISL_495978, EPI_ISL_495979, EPI_ISL_495980, EPI_ISL_495981, EPI_ISL_495982, EPI_ISL_495983, EPI_ISL_495984, EPI_ISL_495985, EPI_ISL_495986, EPI_ISL_495987, EPI_ISL_495988, EPI_ISL_495989, EPI_ISL_495990, EPI_ISL_495991, EPI_ISL_495992, EPI_ISL_495993, EPI_ISL_495994, EPI_ISL_495995, EPI_ISL_495996, EPI_ISL_495997, EPI_ISL_495998, EPI_ISL_495999, EPI_ISL_496000, EPI_ISL_496001, EPI_ISL_496002, EPI_ISL_496003, EPI_ISL_496004, EPI_ISL_496005, EPI_ISL_496006, EPI_ISL_496007, EPI_ISL_496008, EPI_ISL_496009, EPI_ISL_496010, EPI_ISL_496011, EPI_ISL_496012, EPI_ISL_496013, EPI_ISL_496014, EPI_ISL_496015, EPI_ISL_496016, EPI_ISL_496017, EPI_ISL_496018, EPI_ISL_496019, EPI_ISL_496020, EPI_ISL_496021, EPI_ISL_496022, EPI_ISL_496023, EPI_ISL_496024, EPI_ISL_496025, EPI_ISL_496026, EPI_ISL_496027, EPI_ISL_496028, EPI_ISL_496029, EPI_ISL_496030, EPI_ISL_496031, EPI_ISL_496032, EPI_ISL_496033, EPI_ISL_496034, EPI_ISL_496035, EPI_ISL_496036, EPI_ISL_496037, EPI_ISL_496038, EPI_ISL_496039, EPI_ISL_496040, EPI_ISL_496041, EPI_ISL_496042, EPI_ISL_496043, EPI_ISL_496044, EPI_ISL_496045, EPI_ISL_496046, EPI_ISL_496047, EPI_ISL_496048, EPI_ISL_496049, EPI_ISL_496050, EPI_ISL_496051, EPI_ISL_496052, EPI_ISL_496053, EPI_ISL_496054, EPI_ISL_496055, EPI_ISL_496056, EPI_ISL_496057, EPI_ISL_496058, EPI_ISL_496059, EPI_ISL_496060, EPI_ISL_496061, EPI_ISL_496062, EPI_ISL_496063, EPI_ISL_496064, EPI_ISL_496065, EPI_ISL_496066, EPI_ISL_496067, EPI_ISL_496068, EPI_ISL_496069, EPI_ISL_496070, EPI_ISL_496071, EPI_ISL_496072, EPI_ISL_496073, EPI_ISL_496074, EPI_ISL_496075, EPI_ISL_496076, EPI_ISL_496077, EPI_ISL_496078, EPI_ISL_496079, EPI_ISL_496080, EPI_ISL_496081, EPI_ISL_496082, EPI_ISL_496083, EPI_ISL_496084, EPI_ISL_496085, EPI_ISL_496086, EPI_ISL_496087, EPI_ISL_496088, EPI_ISL_496089, EPI_ISL_496090, EPI_ISL_496091, EPI_ISL_496092, EPI_ISL_496093, EPI_ISL_496094, EPI_ISL_496095, EPI_ISL_496096, EPI_ISL_496097, EPI_ISL_496098, EPI_ISL_496099, EPI_ISL_496100, EPI_ISL_496101, EPI_ISL_496102, EPI_ISL_496103, EPI_ISL_496104, EPI_ISL_496105, EPI_ISL_496106, EPI_ISL_496107, EPI_ISL_496108, EPI_ISL_496109, EPI_ISL_496110, EPI_ISL_496111, EPI_ISL_496112, EPI_ISL_496113, EPI_ISL_496114, EPI_ISL_496115, EPI_ISL_496116, EPI_ISL_496117, EPI_ISL_496118, EPI_ISL_496119, EPI_ISL_496120, EPI_ISL_496121, EPI_ISL_496122, EPI_ISL_496123, EPI_ISL_496124, EPI_ISL_496125, EPI_ISL_496126, EPI_ISL_496127, EPI_ISL_496128, EPI_ISL_496129, EPI_ISL_496130, EPI_ISL_496131, EPI_ISL_496132, EPI_ISL_496133, EPI_ISL_496134, EPI_ISL_496135, EPI_ISL_496136, EPI_ISL_496137, EPI_ISL_496138, EPI_ISL_496139, EPI_ISL_496140, EPI_ISL_496141, EPI_ISL_496142, EPI_ISL_496143, EPI_ISL_496144, EPI_ISL_496145, EPI_ISL_496146, EPI_ISL_496147, EPI_ISL_496148, EPI_ISL_496149, EPI_ISL_496150, EPI_ISL_496151, EPI_ISL_496152, EPI_ISL_496153, EPI_ISL_496154, EPI_ISL_496155, EPI_ISL_496156, EPI_ISL_496157, EPI_ISL_496158, EPI_ISL_496159, EPI_ISL_496160, EPI_ISL_496161, EPI_ISL_496162, EPI_ISL_496163, EPI_ISL_496164, EPI_ISL_496165, EPI_ISL_496166, EPI_ISL_496167, EPI_ISL_496168, EPI_ISL_496169, EPI_ISL_496170, EPI_ISL_496171, EPI_ISL_496172, EPI_ISL_496173, EPI_ISL_496174, EPI_ISL_496175, EPI_ISL_496176, EPI_ISL_496177, EPI_ISL_496178, EPI_ISL_496179, EPI_ISL_496180, EPI_ISL_496181, EPI_ISL_496182, EPI_ISL_496183, EPI_ISL_496184, EPI_ISL_496185, EPI_ISL_496186, EPI_ISL_496187, EPI_ISL_496188, EPI_ISL_496189, EPI_ISL_496190, EPI_ISL_496191, EPI_ISL_496192, EPI_ISL_496193, EPI_ISL_496194, EPI_ISL_496195, EPI_ISL_496196, EPI_ISL_496197, EPI_ISL_496198, EPI_ISL_496199, EPI_ISL_496200, EPI_ISL_496201, EPI_ISL_496202, EPI_ISL_496203, EPI_ISL_496204, EPI_ISL_496205, EPI_ISL_496206, EPI_ISL_496207, EPI_ISL_496208, EPI_ISL_496209, EPI_ISL_496210, EPI_ISL_496211, EPI_ISL_496212, EPI_ISL_496213, EPI_ISL_496214, EPI_ISL_496215, EPI_ISL_496216, EPI_ISL_496217, EPI_ISL_496218, EPI_ISL_496219, EPI_ISL_496220, EPI_ISL_496221, EPI_ISL_496222, EPI_ISL_496223, EPI_ISL_496224, EPI_ISL_496225, EPI_ISL_496226, EPI_ISL_496227, EPI_ISL_496228, EPI_ISL_496229, EPI_ISL_496230, EPI_ISL_496231, EPI_ISL_496232, EPI_ISL_496233, EPI_ISL_496234, EPI_ISL_496235, EPI_ISL_496236, EPI_ISL_496237, EPI_ISL_496238, EPI_ISL_496239, EPI_ISL_496240, EPI_ISL_496241, EPI_ISL_496242, EPI_ISL_496243, EPI_ISL_496244, EPI_ISL_496245, EPI_ISL_496246, EPI_ISL_496247, EPI_ISL_496248, EPI_ISL_496249, EPI_ISL_496250, EPI_ISL_496251, EPI_ISL_496252, EPI_ISL_496253, EPI_ISL_496254, EPI_ISL_496255, EPI_ISL_496256, EPI_ISL_496257, EPI_ISL_496258, EPI_ISL_496259, EPI_ISL_496260, EPI_ISL_496261, EPI_ISL_496262, EPI_ISL_496263, EPI_ISL_496264, EPI_ISL_496265, EPI_ISL_496266, EPI_ISL_496267, EPI_ISL_496268, EPI_ISL_496269, EPI_ISL_496270, EPI_ISL_496271, EPI_ISL_496272, EPI_ISL_496273, EPI_ISL_496274, EPI_ISL_496275, EPI_ISL_496276, EPI_ISL_496277, EPI_ISL_496278, EPI_ISL_496279, EPI_ISL_496280, EPI_ISL_496281, EPI_ISL_496282, EPI_ISL_496283, EPI_ISL_496284, EPI_ISL_496285, EPI_ISL_496286, EPI_ISL_496287, EPI_ISL_496288, EPI_ISL_496289, EPI_ISL_496290, EPI_ISL_496291, EPI_ISL_496292, EPI_ISL_496293, EPI_ISL_496294, EPI_ISL_496295, EPI_ISL_496296, EPI_ISL_496297, EPI_ISL_496298, EPI_ISL_496299, EPI_ISL_496300, EPI_ISL_496301, EPI_ISL_496302, EPI_ISL_496303, EPI_ISL_496304, EPI_ISL_496305, EPI_ISL_496306, EPI_ISL_496307, EPI_ISL_496308, EPI_ISL_496309, EPI_ISL_496310, EPI_ISL_496311, EPI_ISL_496312, EPI_ISL_496313, EPI_ISL_496314, EPI_ISL_496315, EPI_ISL_496316, EPI_ISL_496317, EPI_ISL_496318, EPI_ISL_496319, EPI_ISL_496320, EPI_ISL_496321, EPI_ISL_496322, EPI_ISL_496323, EPI_ISL_496324, EPI_ISL_496325, EPI_ISL_496326, EPI_ISL_496327, EPI_ISL_496328, EPI_ISL_496329, EPI_ISL_496330, EPI_ISL_496331, EPI_ISL_496332, EPI_ISL_496333, EPI_ISL_496334, EPI_ISL_496335, EPI_ISL_496336, EPI_ISL_496337, EPI_ISL_496338, EPI_ISL_496931, EPI_ISL_496932, EPI_ISL_496933, EPI_ISL_496934, EPI_ISL_496935, EPI_ISL_496936, EPI_ISL_496937, EPI_ISL_496938, EPI_ISL_496939, EPI_ISL_496940, EPI_ISL_496941, EPI_ISL_496942, EPI_ISL_496943, EPI_ISL_496944, EPI_ISL_496945, EPI_ISL_496946, EPI_ISL_496947, EPI_ISL_496948, EPI_ISL_496949, EPI_ISL_496950, EPI_ISL_496951, EPI_ISL_496952, EPI_ISL_496953, EPI_ISL_496954, EPI_ISL_496955, EPI_ISL_496956, EPI_ISL_496957, EPI_ISL_496958, EPI_ISL_496959, EPI_ISL_496960, EPI_ISL_496961, EPI_ISL_496962, EPI_ISL_496963, EPI_ISL_496964, EPI_ISL_496965, EPI_ISL_496966, EPI_ISL_496967, EPI_ISL_496968, EPI_ISL_496969, EPI_ISL_496970, EPI_ISL_496971, EPI_ISL_496972, EPI_ISL_496973, EPI_ISL_496974, EPI_ISL_496975, EPI_ISL_496976, EPI_ISL_496977, EPI_ISL_496978, EPI_ISL_496979, EPI_ISL_496980, EPI_ISL_496981, EPI_ISL_496982, EPI_ISL_496983, EPI_ISL_496984, EPI_ISL_496985, EPI_ISL_496986, EPI_ISL_496987, EPI_ISL_496988, EPI_ISL_496989, EPI_ISL_496990, EPI_ISL_496991, EPI_ISL_496992, EPI_ISL_496993, EPI_ISL_496994, EPI_ISL_496995, EPI_ISL_496996, EPI_ISL_496997, EPI_ISL_496998, EPI_ISL_496999, EPI_ISL_497000, EPI_ISL_497001, EPI_ISL_497002, EPI_ISL_497003, EPI_ISL_497004, EPI_ISL_497005, EPI_ISL_497006, EPI_ISL_497007, EPI_ISL_497008, EPI_ISL_497009, EPI_ISL_497010, EPI_ISL_497011, EPI_ISL_497012, EPI_ISL_497013, EPI_ISL_497014, EPI_ISL_497015, EPI_ISL_497016, EPI_ISL_497017, EPI_ISL_497018, EPI_ISL_497019, EPI_ISL_497020, EPI_ISL_497021, EPI_ISL_497022, EPI_ISL_497023, EPI_ISL_497024, EPI_ISL_497025, EPI_ISL_497026, EPI_ISL_497027, EPI_ISL_497028, EPI_ISL_497029, EPI_ISL_497030, EPI_ISL_497031, EPI_ISL_497032, EPI_ISL_497033, EPI_ISL_497034, EPI_ISL_497035, EPI_ISL_497036, EPI_ISL_497037, EPI_ISL_497038, EPI_ISL_497039, EPI_ISL_497040, EPI_ISL_497041, EPI_ISL_497042, EPI_ISL_497043, EPI_ISL_497044, EPI_ISL_497045, EPI_ISL_497046, EPI_ISL_497047, EPI_ISL_497048, EPI_ISL_497049, EPI_ISL_497050, EPI_ISL_497051, EPI_ISL_497052, EPI_ISL_497053, EPI_ISL_497054, EPI_ISL_497055, EPI_ISL_497056, EPI_ISL_497057, EPI_ISL_497058, EPI_ISL_497059, EPI_ISL_497060, EPI_ISL_497061, EPI_ISL_497062, EPI_ISL_497063, EPI_ISL_497064, EPI_ISL_497065, EPI_ISL_497066, EPI_ISL_497067, EPI_ISL_497068, EPI_ISL_497069, EPI_ISL_497070, EPI_ISL_497071, EPI_ISL_497072, EPI_ISL_497073, EPI_ISL_497074, EPI_ISL_497075, EPI_ISL_497076, EPI_ISL_497077, EPI_ISL_497078, EPI_ISL_497079, EPI_ISL_497080, EPI_ISL_497081, EPI_ISL_497082, EPI_ISL_497083, EPI_ISL_497084, EPI_ISL_497085, EPI_ISL_497086, EPI_ISL_497087, EPI_ISL_497088, EPI_ISL_497089, EPI_ISL_497090, EPI_ISL_497091, EPI_ISL_497092, EPI_ISL_497093, EPI_ISL_497094, EPI_ISL_497095, EPI_ISL_497096, EPI_ISL_497097, EPI_ISL_497098, EPI_ISL_497099, EPI_ISL_497100, EPI_ISL_497101, EPI_ISL_497102, EPI_ISL_497103, EPI_ISL_497104, EPI_ISL_497105, EPI_ISL_497106, EPI_ISL_497107, EPI_ISL_497108, EPI_ISL_497109, EPI_ISL_497110, EPI_ISL_497111, EPI_ISL_497112, EPI_ISL_497113, EPI_ISL_497114, EPI_ISL_497115, EPI_ISL_497116, EPI_ISL_497117, EPI_ISL_497118, EPI_ISL_497119, EPI_ISL_497120, EPI_ISL_497121, EPI_ISL_497122, EPI_ISL_497123, EPI_ISL_497124, EPI_ISL_497125, EPI_ISL_497126, EPI_ISL_497127, EPI_ISL_497128, EPI_ISL_497129, EPI_ISL_497130, EPI_ISL_497131, EPI_ISL_497132, EPI_ISL_497133, EPI_ISL_497134, EPI_ISL_497135, EPI_ISL_497136, EPI_ISL_497137, EPI_ISL_497138, EPI_ISL_497139, EPI_ISL_497140, EPI_ISL_497141, EPI_ISL_497142, EPI_ISL_497143, EPI_ISL_497144, EPI_ISL_497145, EPI_ISL_497146, EPI_ISL_497147, EPI_ISL_497148, EPI_ISL_497149, EPI_ISL_497150, EPI_ISL_497151, EPI_ISL_497152, EPI_ISL_497153, EPI_ISL_497154, EPI_ISL_497155, EPI_ISL_497156, | | | |

| | | | |
|---|---|---|---|
| EPI_ISL_497157, EPI_ISL_497158, EPI_ISL_497159, EPI_ISL_497160, EPI_ISL_497161, EPI_ISL_497162, EPI_ISL_497163, EPI_ISL_497164, EPI_ISL_497165, EPI_ISL_497166, EPI_ISL_497167, EPI_ISL_497168, EPI_ISL_497169, EPI_ISL_497170, EPI_ISL_497171, EPI_ISL_497172, EPI_ISL_497173, EPI_ISL_497174, EPI_ISL_497175, EPI_ISL_497176, EPI_ISL_497177, EPI_ISL_497178, EPI_ISL_497179, EPI_ISL_497180, EPI_ISL_497181, EPI_ISL_497182, EPI_ISL_497183, EPI_ISL_497184, EPI_ISL_497185, EPI_ISL_497186, EPI_ISL_497187, EPI_ISL_497188, EPI_ISL_497189, EPI_ISL_497190, EPI_ISL_497191, EPI_ISL_497192, EPI_ISL_497193, EPI_ISL_497194, EPI_ISL_497195, EPI_ISL_497196, EPI_ISL_497197, EPI_ISL_497198, EPI_ISL_497199, EPI_ISL_497200, EPI_ISL_497201, EPI_ISL_497202, EPI_ISL_497203, EPI_ISL_497204, EPI_ISL_497205, EPI_ISL_497206, EPI_ISL_497207, EPI_ISL_497208, EPI_ISL_497209, EPI_ISL_497210, EPI_ISL_497211, EPI_ISL_497212, EPI_ISL_497213, EPI_ISL_497214, EPI_ISL_497215, EPI_ISL_497216, EPI_ISL_497217, EPI_ISL_497218, EPI_ISL_497219, EPI_ISL_497220, EPI_ISL_497221, EPI_ISL_497222, EPI_ISL_497223, EPI_ISL_497224, EPI_ISL_497225, EPI_ISL_497226, EPI_ISL_497227, EPI_ISL_497228, EPI_ISL_497229, EPI_ISL_497230, EPI_ISL_497231, EPI_ISL_497232, EPI_ISL_497233, EPI_ISL_497234, EPI_ISL_497235, EPI_ISL_497236, EPI_ISL_497237, EPI_ISL_497238, EPI_ISL_497239, EPI_ISL_497240, EPI_ISL_497241, EPI_ISL_497242, EPI_ISL_497243, EPI_ISL_497244, EPI_ISL_497245, EPI_ISL_497246, EPI_ISL_497247, EPI_ISL_497248, EPI_ISL_497249, EPI_ISL_497250, EPI_ISL_497251, EPI_ISL_497252, EPI_ISL_497253, EPI_ISL_497254, EPI_ISL_497255, EPI_ISL_497256, EPI_ISL_497257, EPI_ISL_497258, EPI_ISL_497259, EPI_ISL_497260, EPI_ISL_497261, EPI_ISL_497262, EPI_ISL_497263, EPI_ISL_497264, EPI_ISL_497265, EPI_ISL_497266, EPI_ISL_497267, EPI_ISL_497268, EPI_ISL_497269, EPI_ISL_497270, EPI_ISL_497271, EPI_ISL_497272, EPI_ISL_497273, EPI_ISL_497274, EPI_ISL_497275, EPI_ISL_497276, EPI_ISL_497277, EPI_ISL_497278, EPI_ISL_497279, EPI_ISL_497280, EPI_ISL_497281, EPI_ISL_497282, EPI_ISL_497283, EPI_ISL_497284, EPI_ISL_497285, EPI_ISL_497286, EPI_ISL_497287, EPI_ISL_497288, EPI_ISL_497289, EPI_ISL_497290, EPI_ISL_497291, EPI_ISL_497292, EPI_ISL_497293, EPI_ISL_497294, EPI_ISL_497295, EPI_ISL_497296, EPI_ISL_497297, EPI_ISL_497298, EPI_ISL_497299, EPI_ISL_497300, EPI_ISL_497301, EPI_ISL_497302, EPI_ISL_497303, EPI_ISL_497304, EPI_ISL_497305, EPI_ISL_497306, EPI_ISL_497307, EPI_ISL_497308, EPI_ISL_497309, EPI_ISL_497310, EPI_ISL_497311, EPI_ISL_497312, EPI_ISL_497313, EPI_ISL_497314, EPI_ISL_497315, EPI_ISL_497316, EPI_ISL_497317, EPI_ISL_497318, EPI_ISL_497319, EPI_ISL_497320, EPI_ISL_497321, EPI_ISL_497322, EPI_ISL_497323, EPI_ISL_497324, EPI_ISL_497325, EPI_ISL_497326, EPI_ISL_497327, EPI_ISL_497328, EPI_ISL_497329, EPI_ISL_497330, EPI_ISL_497331, EPI_ISL_497332, EPI_ISL_497333, EPI_ISL_497334, EPI_ISL_497335, EPI_ISL_497336, EPI_ISL_497337, EPI_ISL_497338, EPI_ISL_497339, EPI_ISL_497340, EPI_ISL_497342, EPI_ISL_497343, EPI_ISL_497344, EPI_ISL_497345, EPI_ISL_497346, EPI_ISL_497347, EPI_ISL_497348, EPI_ISL_497349, EPI_ISL_497350, EPI_ISL_497351, EPI_ISL_497352, EPI_ISL_497353, EPI_ISL_497354, EPI_ISL_497355, EPI_ISL_497356, EPI_ISL_497357, EPI_ISL_497358, EPI_ISL_497359, EPI_ISL_497360, EPI_ISL_497361, EPI_ISL_497362, EPI_ISL_497363, EPI_ISL_497364, EPI_ISL_497365, EPI_ISL_497366, EPI_ISL_497367, EPI_ISL_497368, EPI_ISL_497369, EPI_ISL_497370, EPI_ISL_497371, EPI_ISL_497372, EPI_ISL_497373, EPI_ISL_497374, EPI_ISL_497375, EPI_ISL_497376, EPI_ISL_497377, EPI_ISL_497378, EPI_ISL_497379, EPI_ISL_497380, EPI_ISL_497381, EPI_ISL_497382, EPI_ISL_497383, EPI_ISL_497384, EPI_ISL_497385, EPI_ISL_497386, EPI_ISL_497387, EPI_ISL_497388, EPI_ISL_497389, EPI_ISL_497390, EPI_ISL_497391, EPI_ISL_497392, EPI_ISL_497393, EPI_ISL_497394, EPI_ISL_497395, EPI_ISL_497396, EPI_ISL_497397, EPI_ISL_497398, EPI_ISL_497399, EPI_ISL_497400, EPI_ISL_497401, EPI_ISL_497402, EPI_ISL_497403, EPI_ISL_497404, EPI_ISL_497405, EPI_ISL_497406, EPI_ISL_497407, EPI_ISL_497408, EPI_ISL_497409, EPI_ISL_497410, EPI_ISL_497411, EPI_ISL_497412, EPI_ISL_497413, EPI_ISL_497414, EPI_ISL_497415, EPI_ISL_497416, EPI_ISL_497417, EPI_ISL_497418, EPI_ISL_497419, EPI_ISL_497420, EPI_ISL_497421, EPI_ISL_497422, EPI_ISL_497423, EPI_ISL_497424, EPI_ISL_497425, EPI_ISL_497426, EPI_ISL_497427, EPI_ISL_497428, EPI_ISL_497429, EPI_ISL_497430, EPI_ISL_497431, EPI_ISL_497432, EPI_ISL_497433, EPI_ISL_497434, EPI_ISL_497435, EPI_ISL_497436, EPI_ISL_497437, EPI_ISL_497438, EPI_ISL_497439, EPI_ISL_497440, EPI_ISL_497441, EPI_ISL_497442, EPI_ISL_497443, EPI_ISL_497444, EPI_ISL_497445, EPI_ISL_497446, EPI_ISL_497447, EPI_ISL_497448, EPI_ISL_497449, EPI_ISL_497450, EPI_ISL_497451, EPI_ISL_497452, EPI_ISL_497453, EPI_ISL_497454, EPI_ISL_497455, EPI_ISL_497456, EPI_ISL_497457, EPI_ISL_497458, EPI_ISL_497459, EPI_ISL_497460, EPI_ISL_497461, EPI_ISL_497462, EPI_ISL_497463, EPI_ISL_497464, EPI_ISL_497465, EPI_ISL_497466, EPI_ISL_497467, EPI_ISL_497468, EPI_ISL_497469, EPI_ISL_497470, EPI_ISL_497471, EPI_ISL_497472, EPI_ISL_497473, EPI_ISL_497474, EPI_ISL_497475, EPI_ISL_497476, EPI_ISL_497477, EPI_ISL_497478, EPI_ISL_497479, EPI_ISL_497480, EPI_ISL_497481, EPI_ISL_497482, EPI_ISL_497483, EPI_ISL_497484, EPI_ISL_497485, EPI_ISL_497486, EPI_ISL_497487, EPI_ISL_497488, EPI_ISL_497489, EPI_ISL_497490, EPI_ISL_497491, EPI_ISL_497492, EPI_ISL_497493, EPI_ISL_497494, EPI_ISL_497495, EPI_ISL_497496, EPI_ISL_497497, EPI_ISL_497498, EPI_ISL_497499, EPI_ISL_497500, EPI_ISL_497501, EPI_ISL_497502, EPI_ISL_497503, EPI_ISL_497504, EPI_ISL_497505, EPI_ISL_497506, EPI_ISL_497507, EPI_ISL_497508, EPI_ISL_497509, EPI_ISL_497510, EPI_ISL_497511, EPI_ISL_497512, EPI_ISL_497513, EPI_ISL_497514, EPI_ISL_497515, EPI_ISL_497516, EPI_ISL_497517, EPI_ISL_497518, EPI_ISL_497519, EPI_ISL_497520, EPI_ISL_497521, EPI_ISL_497522, EPI_ISL_497523, EPI_ISL_497524, EPI_ISL_497525, EPI_ISL_497526, EPI_ISL_497527, EPI_ISL_497528, EPI_ISL_497529, EPI_ISL_497530, EPI_ISL_497531, EPI_ISL_497532, EPI_ISL_497533, EPI_ISL_497534, EPI_ISL_497535, EPI_ISL_497536, EPI_ISL_497537, EPI_ISL_497538, EPI_ISL_497539, EPI_ISL_497540, EPI_ISL_497541, EPI_ISL_497542, EPI_ISL_497543, EPI_ISL_497544, EPI_ISL_497545, EPI_ISL_497546, EPI_ISL_497547, EPI_ISL_497548, EPI_ISL_497549, EPI_ISL_497550, EPI_ISL_497551, EPI_ISL_497552, EPI_ISL_497553, EPI_ISL_497554, EPI_ISL_497555, EPI_ISL_497556, EPI_ISL_497557, EPI_ISL_497558, EPI_ISL_497559, EPI_ISL_497560, EPI_ISL_497561, EPI_ISL_497562, EPI_ISL_497563, EPI_ISL_497564, EPI_ISL_497565, EPI_ISL_497566, EPI_ISL_497567, EPI_ISL_497568, EPI_ISL_497569, EPI_ISL_497570, EPI_ISL_497571, EPI_ISL_497572, EPI_ISL_497573, EPI_ISL_497574, EPI_ISL_497575, EPI_ISL_497576, EPI_ISL_497577, EPI_ISL_497578, EPI_ISL_497579, EPI_ISL_497580, EPI_ISL_497581, EPI_ISL_497582, EPI_ISL_497583, EPI_ISL_497584, EPI_ISL_497585, EPI_ISL_497586, EPI_ISL_497587, EPI_ISL_497588, EPI_ISL_497589, EPI_ISL_497590, EPI_ISL_497591, EPI_ISL_497592, EPI_ISL_497593, EPI_ISL_497594, EPI_ISL_497595, EPI_ISL_497596, EPI_ISL_497597, EPI_ISL_497598, EPI_ISL_497599, EPI_ISL_497600, EPI_ISL_497601, EPI_ISL_497602, EPI_ISL_497603, EPI_ISL_497604, EPI_ISL_497605, EPI_ISL_497606, EPI_ISL_497607, EPI_ISL_497608, EPI_ISL_497609, EPI_ISL_497610, EPI_ISL_497611, EPI_ISL_497612, EPI_ISL_497613, EPI_ISL_497614, EPI_ISL_497615, EPI_ISL_497616, EPI_ISL_497617, EPI_ISL_497618, EPI_ISL_497619, EPI_ISL_497620, EPI_ISL_497621, EPI_ISL_497622, EPI_ISL_497623, EPI_ISL_497624, EPI_ISL_497625, EPI_ISL_497626, EPI_ISL_497627, EPI_ISL_497628, EPI_ISL_497629, EPI_ISL_497630, EPI_ISL_497631, EPI_ISL_497632, EPI_ISL_497633, EPI_ISL_497634, EPI_ISL_497635, EPI_ISL_497636, EPI_ISL_497637, EPI_ISL_497638, EPI_ISL_497639, EPI_ISL_497640, EPI_ISL_497641, EPI_ISL_497642, EPI_ISL_497643, EPI_ISL_497644, EPI_ISL_497645, EPI_ISL_497646, EPI_ISL_497647, EPI_ISL_497648, EPI_ISL_497649, EPI_ISL_497650, EPI_ISL_497651, EPI_ISL_497652, EPI_ISL_497653, EPI_ISL_497654, EPI_ISL_497655, EPI_ISL_497656, EPI_ISL_497657, EPI_ISL_497658, EPI_ISL_497659, EPI_ISL_497660, EPI_ISL_497661, EPI_ISL_497662, EPI_ISL_497663, EPI_ISL_497664, EPI_ISL_497665, EPI_ISL_497666, EPI_ISL_497667, EPI_ISL_497668, EPI_ISL_497669, EPI_ISL_497670, EPI_ISL_497671, EPI_ISL_497672, EPI_ISL_497673, EPI_ISL_497674, EPI_ISL_497675, EPI_ISL_497676, EPI_ISL_497677, EPI_ISL_497678, EPI_ISL_497679, EPI_ISL_497680, EPI_ISL_497681, EPI_ISL_497682, EPI_ISL_497683, EPI_ISL_497684, EPI_ISL_497685, EPI_ISL_497686, EPI_ISL_497687, EPI_ISL_497688, EPI_ISL_497689, EPI_ISL_497690, EPI_ISL_497691, EPI_ISL_497692, EPI_ISL_497693, EPI_ISL_497694, EPI_ISL_497695, EPI_ISL_497696, EPI_ISL_497697, EPI_ISL_497698, EPI_ISL_497699, EPI_ISL_497700, EPI_ISL_497701, EPI_ISL_497702, EPI_ISL_497703, EPI_ISL_497704, EPI_ISL_497705, EPI_ISL_497706, EPI_ISL_497707, EPI_ISL_497708, EPI_ISL_497709, EPI_ISL_497710, EPI_ISL_497711, EPI_ISL_497712, EPI_ISL_497713, EPI_ISL_497714, EPI_ISL_497715, EPI_ISL_497716, EPI_ISL_497717, EPI_ISL_497718, EPI_ISL_497719, EPI_ISL_497720, EPI_ISL_497721, EPI_ISL_497722, EPI_ISL_497723, EPI_ISL_497724, EPI_ISL_497725, EPI_ISL_497726 | | | |
| see above | Washington State Department of Health | Seattle Flu Study | Deborah A. Nickerson, Chris D. Frazar, Jover Lee, Benjamin Pelle, Matthew Richardson, Amanda Adler, Elisabeth Brandstetter, Peter D. Han, Kairsten Fay, Misja Ilcisin, Kirsten Lacombe, Thomas R. Sibley, Melissa Truong, Caitlin R. Wolf, Romesh Gautom, Geoff Melly, Brian Hiatt, Philip Dykema, Scott Lindquist, Michael Boeckh, Janet A. Englund, Michael Famulare, Barry R. Lutz, Mark J. Rieder, Lea M. Starita, Matthew Thompson, Helen Y. Chu, Jay Shendure, Trevor Bedford |
| EPI_ISL_497872 | UW Virology Lab | UW Virology Lab | Pavitra Roychoudhury, Amin Addetia, Hong Xie, Lasata Shrestha, Truong Nguyen, Meei-Li Huang, Keith Jerome, Alexander Greninger |
| EPI_ISL_498744, EPI_ISL_498745, EPI_ISL_498746, EPI_ISL_498747 | Quest Diagnostics | Quest Diagnostics | Rosenthal,S.H., Gerasimova,A., Kagan,R.M. and Owen, R. |
| EPI_ISL_500512 | Mayo Clinic Laboratories | University of Washington Virology Lab | Pavitra Roychoudhury, Hong Xie, Lasata Shrestha, Amin Addetia, Truong Nguyen, Victoria M Rachleff, Meei-Li Huang, Keith R Jerome, Alexander Greninger |
| EPI_ISL_501086, EPI_ISL_501087, EPI_ISL_501088, EPI_ISL_501089, EPI_ISL_501090, EPI_ISL_501091, EPI_ISL_501092, EPI_ISL_501093, EPI_ISL_501094, EPI_ISL_501095, EPI_ISL_501096, EPI_ISL_501097, EPI_ISL_501098, EPI_ISL_501099, EPI_ISL_501100, EPI_ISL_501101, EPI_ISL_501102, EPI_ISL_501103, EPI_ISL_501104, EPI_ISL_501105, EPI_ISL_501106, EPI_ISL_501107, EPI_ISL_501108, EPI_ISL_501109, EPI_ISL_501110, EPI_ISL_501111, EPI_ISL_501112, EPI_ISL_501113, EPI_ISL_501114, EPI_ISL_501115, EPI_ISL_501116, EPI_ISL_501117, EPI_ISL_501119, EPI_ISL_501121, EPI_ISL_501122, EPI_ISL_501123, EPI_ISL_501125, EPI_ISL_501126, EPI_ISL_501127, EPI_ISL_501128, EPI_ISL_501151, EPI_ISL_501152, EPI_ISL_501153, EPI_ISL_501154, EPI_ISL_501155, EPI_ISL_501156, EPI_ISL_501157, EPI_ISL_501158, EPI_ISL_501159, EPI_ISL_501160, EPI_ISL_501161, EPI_ISL_501162, EPI_ISL_501163, EPI_ISL_501164 | | | |
| see above | University of Washington Virology Lab | University of Washington Virology Lab | Pavitra Roychoudhury, Hong Xie, Lasata Shrestha, Amin Addetia, Truong Nguyen, Victoria M Rachleff, Meei-Li Huang, Keith R Jerome, Alexander Greninger |
| EPI_ISL_509059, EPI_ISL_509062, EPI_ISL_509063, EPI_ISL_509064, EPI_ISL_509065, EPI_ISL_509066, EPI_ISL_509067, EPI_ISL_509068, EPI_ISL_509078, EPI_ISL_509079, EPI_ISL_509080, EPI_ISL_509081, EPI_ISL_509082, EPI_ISL_509083, EPI_ISL_509084, EPI_ISL_509085, EPI_ISL_509088, EPI_ISL_509089, EPI_ISL_509090, EPI_ISL_509092, EPI_ISL_509093, EPI_ISL_509094, EPI_ISL_509095, EPI_ISL_509096, EPI_ISL_509097, EPI_ISL_509100, EPI_ISL_509102, EPI_ISL_509103, EPI_ISL_509106, EPI_ISL_509111, EPI_ISL_509112, EPI_ISL_509116, EPI_ISL_509117, EPI_ISL_509118, EPI_ISL_509119, EPI_ISL_509121, EPI_ISL_509122, EPI_ISL_509123, EPI_ISL_509124, EPI_ISL_509125, EPI_ISL_509134, EPI_ISL_509135, EPI_ISL_509136, EPI_ISL_509137, EPI_ISL_509138 | | | |
| see above | OHSU Lab Services Molecular Microbiology Lab | Oregon SARS-CoV-2 Genome Sequencing Center | Brendan L. O'Connell, Ruth V. Nichols, Sally B. Grindstaff, Alec J. Hirsch, Guang Fan, Daniel N. Streblow, William B. Messer, Andrew C. Adey, Benjamin N. Bimber, Brian J. O'Roak |
| EPI_ISL_509829, EPI_ISL_509835, EPI_ISL_509838, EPI_ISL_509840, EPI_ISL_509842, EPI_ISL_509940, EPI_ISL_509946, EPI_ISL_509947, EPI_ISL_509952, EPI_ISL_509954, EPI_ISL_509957, EPI_ISL_509958, EPI_ISL_509959 | | | |
| see above | University of Wisconsin-Madison AIDS Vaccine Research Laboratories | University of Wisconsin-Madison AIDS Vaccine Research Laboratories | Gage Moreno, Katarina Braun, et al. AIDS Vaccine Research Laboratories |
| EPI_ISL_511852, EPI_ISL_511853, EPI_ISL_511854, EPI_ISL_511855, EPI_ISL_511856, EPI_ISL_511857, EPI_ISL_511858, EPI_ISL_511859, EPI_ISL_511860, EPI_ISL_511861 | UW Virology Lab | UW Virology Lab | Pavitra Roychoudhury, Amin Addetia, Hong Xie, Lasata Shrestha, Truong Nguyen, Meei-Li Huang, Keith Jerome, Alexander Greninger |
| EPI_ISL_511863, EPI_ISL_511864 | Johns Hopkins Hospital Department of Pathology | Johns Hopkins Hospital Department of Pathology | Peter M. Thielen, Thomas Mehoke, Shirlee Wohl, Srividya Ramakrishnan, Melanie Kirsche, Amanda Ernlund, Craig Howser, Kristina Zudock, Oluwaseun Falade-Nwulia, Norah Sadowski, Paul Morris, Mark Hopkins, Yunfan Fan, Nidia Trovao, Victoria Gniazdowski, Michael C. Schatz, Stuart C. Ray, Winston Timp, Heba H. Mostafa |
| EPI_ISL_512086 | UW Virology lab | UW Virology lab | Pavitra Roychoudhury, Amin Addetia, Hong Xie, Lasata Shrestha, Truong Nguyen, Meei-Li Huang, Keith Jerome, Alexander Greninger |
| EPI_ISL_513632, EPI_ISL_513633, EPI_ISL_513634, EPI_ISL_513635, EPI_ISL_513636, EPI_ISL_513637, EPI_ISL_513638, EPI_ISL_515271, EPI_ISL_515272, EPI_ISL_515273, EPI_ISL_515274, EPI_ISL_515275, EPI_ISL_515276, EPI_ISL_515277, EPI_ISL_515278, EPI_ISL_515279, EPI_ISL_515280, EPI_ISL_515281, EPI_ISL_515282, EPI_ISL_515283, EPI_ISL_515284, EPI_ISL_515285, EPI_ISL_515286 | | | |
| see above | University of Washington Virology Lab | University of Washington Virology Lab | Pavitra Roychoudhury, Hong Xie, Lasata Shrestha, Amin Addetia, Truong Nguyen, Victoria M Rachleff, Meei-Li Huang, Keith R Jerome, Alexander Greninger |
